# A Polymorphic Residue That Attenuates the Antiviral Potential of Interferon Lambda 4 in Hominid Lineages

**DOI:** 10.1101/214825

**Authors:** Connor G. G. Bamford, Elihu Aranday-Cortes, Inès Cordeiro Filipe, Swathi Sukumar, Daniel Mair, Ana da Silva Filipe, Juan L. Mendoza, K. Christopher Garcia, Shaohua Fan, Sarah A. Tishkoff, John McLauchlan

**Author notes:** Corresponding Author/Lead Contact: John McLauchlan.

## Abstract

As antimicrobial signalling molecules, type III or lambda interferons (IFNλs) are critical for defence against infection by diverse pathogens. Counter-intuitively, expression of one member of the family, IFNλ4, is associated with decreased clearance of hepatitis C virus (HCV) in the human population; by contrast, a natural in-frame nucleotide insertion that abrogates IFNλ4 production improves viral clearance. To further understand how genetic variation between and within species affects IFNλ4 function, we screened a panel of extant coding variants of human IFNλ4 and identified three variants that substantially affect antiviral activity (P70S, L79F and K154E). The most notable variant was K154E, which enhanced *in vitro* activity in a range of antiviral and interferon stimulated gene (ISG) assays. This more active E154 variant of IFNλ4 was found only in African Congo rainforest ‘Pygmy’ hunter-gatherers. Remarkably, E154 was highly conserved as the ancestral residue in mammalian IFNλ4s yet K154 is the dominant variant throughout evolution of the hominid genus *Homo*. Compared to chimpanzee IFNλ4, the human orthologue had reduced activity due to amino acid substitution of glutamic acid with lysine at position 154. Meta-analysis of published gene expression data from humans and chimpanzees showed that this difference in activity between K154 and E154 in IFNλ4 is consistent with differences in antiviral gene expression *in vivo* during HCV infection. Mechanistically, our data suggest that human-specific K154 likely affects IFNλ4 activity by reducing secretion and potency. We postulate that evolution of an IFNλ4 with attenuated activity in humans (K154) likely contributes to distinct host-specific responses to and outcomes of infection, such as HCV.

## Introduction

Vertebrates have evolved the capacity to coordinate their antiviral defences through the action of proteins called interferons (IFNs) (1), which are small secreted signalling proteins produced by cells after sensing viral infection. IFNs bind to cell surface receptors, commencing autocrine and paracrine signalling via the JAK-STAT pathway. Through this mechanism, IFNs induce expression of hundreds of ‘interferon-stimulated genes’ (ISGs) that establish a cell-intrinsic ‘antiviral state’ and regulate cellular immunity and inflammation (2, 3). Thus, IFNs are pleiotropic in activity and modulate aspects of protective immunity and pathogenesis (4).

Three groups of IFNs have been identified (types I – III), with the type III family (termed IFNλs) being the most recently discovered (5, 6). Emerging evidence highlights the critical and non-redundant role that IFNλs play in protecting against diverse pathogens, including viruses, such as norovirus (7), influenza virus (8) and flaviviruses (9); bacteria (10); and fungi (11). While IFNλs induce nearly identical genes to type I IFNs, differences in signalling kinetics and cell-type specificity contribute to their specialisation (12, 13). Hence, as a consequence of selective expression of the IFNλ receptor 1 (IFNλR1) co-receptor on epithelial cells (13), type III IFNs play a significant role in defence of ‘barrier tissues’, such as the gut, respiratory tract and liver (reviewed in 14); the second co-receptor for IFNλ is IL10-R2, which is expressed more broadly.

Although important for host defence, some IFNs are highly polymorphic (15). In humans, a number of genetic variants in the type III IFN locus (containing IFNλs 1 – 4) have been identified and are associated with clinical phenotypes relating to viral infection (16–18). Although many of these variants are in linkage disequilibrium, the major functional variant is thought to lie in the *IFNL4* gene (19). This causative variant is a single substitution/insertion mutation converting the ‘ ΔG’ allele to a ‘TT’ allele (rs368234815), thereby yielding a frameshift which leads to loss of active human IFNλ4 (HsIFNλ4) (18). Genome-wide association studies have convincingly demonstrated a seemingly counter-intuitive correlation between the *IFNL4* ΔG allele and reduced clearance of hepatitis C virus (HCV) infection, i.e individuals who produce HsIFNλ4 clear HCV infection with reduced frequency in the presence or absence of antiviral IFN therapy (17, 18). Although IFNλ4 is highly conserved among mammals, the ‘pseudogenising’ TT allele of HsIFNλ4 has evolved under positive selection in some human populations suggesting that expression of the wild-type protein likely conferred a fitness cost during recent human evolution (20). Expression of IFNλ4 is tightly controlled and reduced in human as well as Gorilla cells following viral infection compared to IFNλ3 (21). The mechanism underlying the contribution of HsIFNλ4 to viral persistence in HCV infection is not well understood but is associated with enhanced ISG induction. Moreover, a common natural variant of HsIFNλ4 (P70S) (18), which has reduced signalling capacity, is also linked with improved HCV clearance (22). Thus, there is a spectrum of HsIFN□4 activity in humans as a consequence of natural variation that has a significant influence on chronic HCV infection.

Whether other human IFNλ4 variants exist in addition to P70S, which affect antiviral activity, has not been explored fully. In this study, we have examined human genetic data to identify other possible naturally occurring IFNλ4 variants and performed comparative analysis with mammalian orthologues in species closely related to humans. We provide evidence that the antiviral potential for the most common form of IFNλ4 in humans has attenuated activity due to a single amino acid substitution. In addition, we propose that acquisition of the attenuating substitution arose very early during human evolution but that some populations do encode a more active variant. Mechanistically, our data suggest that the reduced antiviral potential of human IFNλ4 results from a likely defect in secretion and potency.

## Results

### Functional consequences of human IFNλ4 non-synonymous variation

Firstly, we undertook genetic and functional comparisons of natural human IFNλ4 coding variants present in the human population. We identified 15 non-synonymous HsIFNλ4 variants in the 1000 Genomes Project database (23) (**Fig 1A and Supplementary Data File 1**), including three previously described variants (C17Y, P60R and P70S; >1% global frequency, classified as ‘common’) (18). The remaining 12 variants were classified as rare (<1% global frequency). The African population harboured the largest number of, as well as the most unique, variants. Interestingly, three rare variants (A8S, S56R and L79F) were shared exclusively between African and American populations, which may have arisen due to relatively recent movements of people perhaps through the transatlantic slave trade.

**Fig 1.**
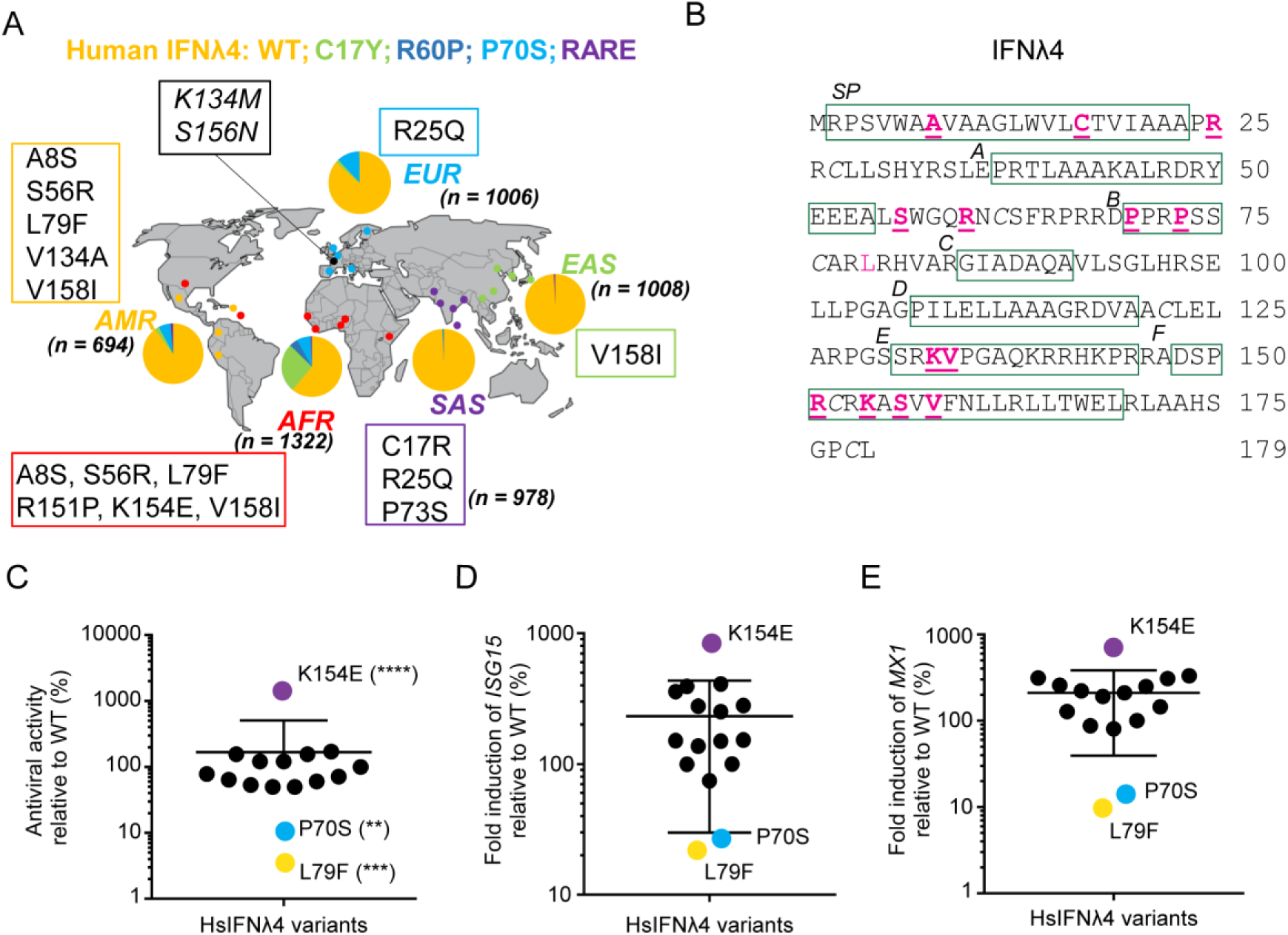
Rare non-synonymous variants of HsIFNλ4 affect antiviral activity. (A) Ancestry-based localization and frequency of human non-synonymous variants of HsIFNλ4 in African (AFR), South Asian (SAS), East Asian (EAS), European (EUR) and American (AMR) populations within the 1000 Genomes dataset. ‘n’ represents the number of alleles tested in each population. Common and rare variants are those which have frequencies of >1% and <1% respectively in the 1000 Genome data. Common variants include: wt (orange), C17Y (light green), R60P (dark blue) and P70S (cyan). Rare variants (purple) include: A8S, C17R, R25Q, S56R, P73S, L79F, K133M, V134A, R151P, K154E, S156N, and V158I. Variants K133M and S156N (black) did not have an associated ethnicity but were found in the dataset from the Netherlands (Genome of the Netherlands cohort) (69). (B) Location of non-synonymous variants in the HsIFNλ4 polypeptide (underlined pink). Regions of predicted structural significance are boxed (green), including the signal peptide (SP) and helices (A to F) (24). There is a single N-linked glycosylation site at position 61 (N61). Note that there are 2 non-synonymous changes at C17 (C17R and C17Y). Cysteine residues involved in disulphide bridge formation are italicised. See Supplementary Data File 1 for genetic identifiers for the variants described here. (C) Antiviral activity of all HsIFNλ4 natural variants in an anti-EMCV CPE assay relative to wt protein in HepaRG cells. Cells were stimulated with serial dilutions of HsIFNλ4-containing CM for 24 hrs and then infected with EMCV (MOI = 0.3 PFU/cell) for 24 hrs at which point CPE was assessed by crystal violet staining. After staining, the dilution providing ∼50% protection was determined. Mean of combined data from three independent experiments (n=3) are shown. Error bars represent mean and SEM for all variants combined. Data are shown in S2 Fig A. **** = <0.0001; *** = <0.001; ** = <0.01 by one-way ANOVA compared to wt with a Dunnett’s test to correct for multiple comparisons. Controls (HsIFNλ4-TT and EGFP) are not shown but gave no protection against EMCV in the assay. Those variants with >2-fold change are highlighted with colours: purple (K154E,), cyan (P70S) and yellow (L79F). (D and E) ISG gene expression determined by RT-qPCR following stimulation of cells with HsIFNλ4 variants. Relative fold change of ISG15 (D) and Mx1 mRNAs (E) in HepaRG cells stimulated with CM (1:4 dilution) from plasmid-transfected cells compared to wt HsIFNλ4. Cells were stimulated for 24 hrs. Data points show mean of biological replicates (n=3) and the error bar represents mean and SEM for all variants combined. Expanded data are shown in S2 Fig B and C. Variants are coloured based on antiviral assays described in Fig 1C. Numerical data used for graph construction are available in **Supplementary Data File 4 sheet 1.**

Variants were located in regions of functional significance in the HsIFNλ4 protein (**Fig 1B and S1 Fig A-C**), such as the predicted signal peptide (amino acids 1-24), surrounding the single glycosylation site (N61) [both of which are required for secretion of active protein], and helix F that is predicted to interact with the IFNλR1 receptor (variants 151–158) (24). Interestingly, the variants in helix F were clustered in the N-terminal portion of the predicted helix. Based on the above predictions, we hypothesised that some of these variants may have phenotypic effects on HsIFNλ4 function. Of note, no variants were found in helix D, which is predicted to contribute to interaction with the IL-10R2 receptor, nor on the IL-10R2-interacting face of the protein.

The functional impact of variation on HsIFNλ4 has only been assessed for the common P70S variant and so we sought to screen all other variants in activity assays. To determine whether variants affected HsIFNλ4 antiviral activity, they were introduced independently into an expression plasmid that produced HsIFNλ4 with a C-terminal ‘FLAG’ tag. Transient transfection of the expression plasmids into human ‘producer’ cells (HEK-293T cells) allowed harvesting of active HsIFNλ4 in the cell supernatant (referred to herein as conditioned media [CM]) thereby enabling analysis of the effects of variants on HsIFNλ4 production, glycosylation, secretion and potency; a similar approach has been successfully adopted previously to determine the relative activities of secreted HsIFNλ3 and HsIFNλ4 as well as a HsIFNλ4 variant that is not glycosylated (24). We chose to screen the function of the panel of HsIFNλ4 variants on the interferon-competent hepatocyte cell line HepaRG cells (25).

Firstly, we investigated the antiviral activity of variants by titrating them against encephalomyocarditis virus (EMCV), a highly IFN-sensitive and cytopathic virus used to measure IFN-mediated protection (26) (**Fig 1C and S2 Fig A**). We also measured their capacity to induce two major ISGs, *MX1* and *ISG15*, by RT-qPCR (**Fig 1D and E, and S2 Fig B and C**) and validated the ISG15 mRNA data by determining production of unconjugated ‘mono’ ISG15 and high-molecular weight ISG15-conjugates (‘ISGylation’; **S2 Fig D**). We also constructed a series of negative controls (plasmids expressing EGFP and the frameshift TT variant of HsIFNλ4), a positive control (HsIFNλ3op) for comparative analysis to examine HsIFNλ4 activity, and three HsIFNλ4 variants, which do not occur naturally but were included as they could alter post-translational modification (N61A which ablates glycosylation) or potential receptor interactions (F159A and L162A located in helix F), respectively (27). Negative controls (EGFP or the frameshift TT variant) gave very low induction of *ISG15* and *MX1* and no detectable antiviral activity in the EMCV assay whereas the positive control (HsIFNλ3op) was highly active in both assays (**S2 Fig A-C**). The non-natural variants N61A and F159A almost abolished activity compared to wt HsIFNλ4 and HsIFNλ3op while L162A gave slightly less activity in the ISG induction assay but activity was reduced to a greater extent in the EMCV assay. In a previous report, ablating glycosylation at N61 substantially reduced activity of secreted HsIFNλ4 in an ISG induction assay (24). Thus, our assay systems recapitulated findings from previous studies with similar assays and provided a range of activities to assess the impact of the natural HsIFNλ4 variants.

Our analyses on the natural HsIFNλ4 variants revealed that only three variants (P70S, L79F and K154E) consistently and substantially modulated antiviral activity and signalling compared to wt HsIFNλ4 (**Fig 1C-E**). The impact of these variants was particularly pronounced in the EMCV assay that measures the dilution giving 50% activity over a large range of dilutions (**Fig 1C**). Our results confirmed previous observations on the lower activity of the P70S variant (22) and demonstrated that the rare L79F variant had a similar phenotype. By contrast, the K154E variant substantially enhanced antiviral activity and ISG induction.

These effects on activity for P70S, L79F and K154E did not arise from differences in the levels of HsIFNλ4 intracellular production or changes to glycosylation (**S3 Fig A and B**). However, variants S56R and R60P (R60P is a common variant in Africa) did lead to marked reductions in the glycosylated form of HsIFNλ4 as demonstrated by the mean ratio of glycosylated:non-glycosylated protein (**S3 Fig**) but did not greatly alter their antiviral activity in contrast with our findings with the N61A non-natural variant, which abolished both glycosylation and antiviral activity of conditioned media (**S2 Fig A-C and S3 Fig A and B**). From this screen, we concluded that three non-synonymous variants in HsIFNλ4 (P70S, L79F and K154E), identified as either common or rare alleles in the human population, affect activity of the protein.

Examining the global distribution of genetic variation can help understand its origins, evolution and functional consequences. P70S is a common variant that is found worldwide (in every population in the 1000 Genomes Database). By contrast, L79F and K154E are rare and geographically restricted to West Africa/Americas, and central Africa, respectively (**Fig 1A**). From further interrogation of human genome datasets (28), the HsIFNλ4 K154E variant was present in two individuals from different African rainforest ‘Pygmy’ hunter-gatherer populations (Baka and Bakola) in Cameroon (**S4 Fig A**). The Bakola individual was homozygous for the ΔG allele, indicating that the K154E variant would be encoded on one of the functional ΔG HsIFN□4 alleles. The Baka subject was heterozygous at rs368234815 (ΔG/TT) and thus could produce either wt or the more active K154E form of HsIFNλ4. Each of these individuals also had additional non-synonymous HsIFNλ4 variants (V158I and R151P, Baka and Bakola individuals respectively); these variants were included in our functional screen of HsIFNλ4 variants but did not significantly alter activity (**Fig 1C-E and S2 Fig A-C**). K154E was not found in other East or Southern African hunter-gatherer populations (such as Hadza and Sandawe) nor in the African San, who have the oldest genetic lineages among humans (29) (**S4 Fig B**); it was also not identified in Neanderthal and Denisovan lineages (denoted as ‘archaic’ in **S4 Fig B**). However, E154 is encoded in the IFNλ4 orthologue for the chimpanzee, *Pan troglodytes* (Pt), our closest mammalian species. Notably, the human TT allele encodes a potential K154 codon (data not shown) suggesting that the E154K substitution arose in humans prior to *IFNL4* pseudogenisation. Together with the fact that nearly all humans encode K154, these data suggest that the less active E154K substitution emerged early during human evolution after the divergence of our last common ancestor with chimpanzees.

### Functional comparison of primate IFNλ4 orthologues

Since a lysine residue encoded at positon 154 is unique to humans compared to other mammalian species (**Fig 2A**) (30), we compared wt HsIFNλ4 and its K154E variant to wt PtIFNλ4 and an equivalent ‘humanised’ PtIFNλ4 E154K mutant in both the EMCV and ISG induction assays as well as a CRISPR-Cas9 cell line in which the EGFP coding region had been introduced into the endogenous *ISG15* gene upstream of and in-frame with the *ISG15* open reading frame (ORF) (**S5 Fig**). This cell line offered advantages over other approaches since it facilitated measurement of ISG induction of an endogenous gene by assessing EGFP fluorescence across a range of dilutions of secreted IFNλs (**S5 Fig B**).

**Fig 2.**
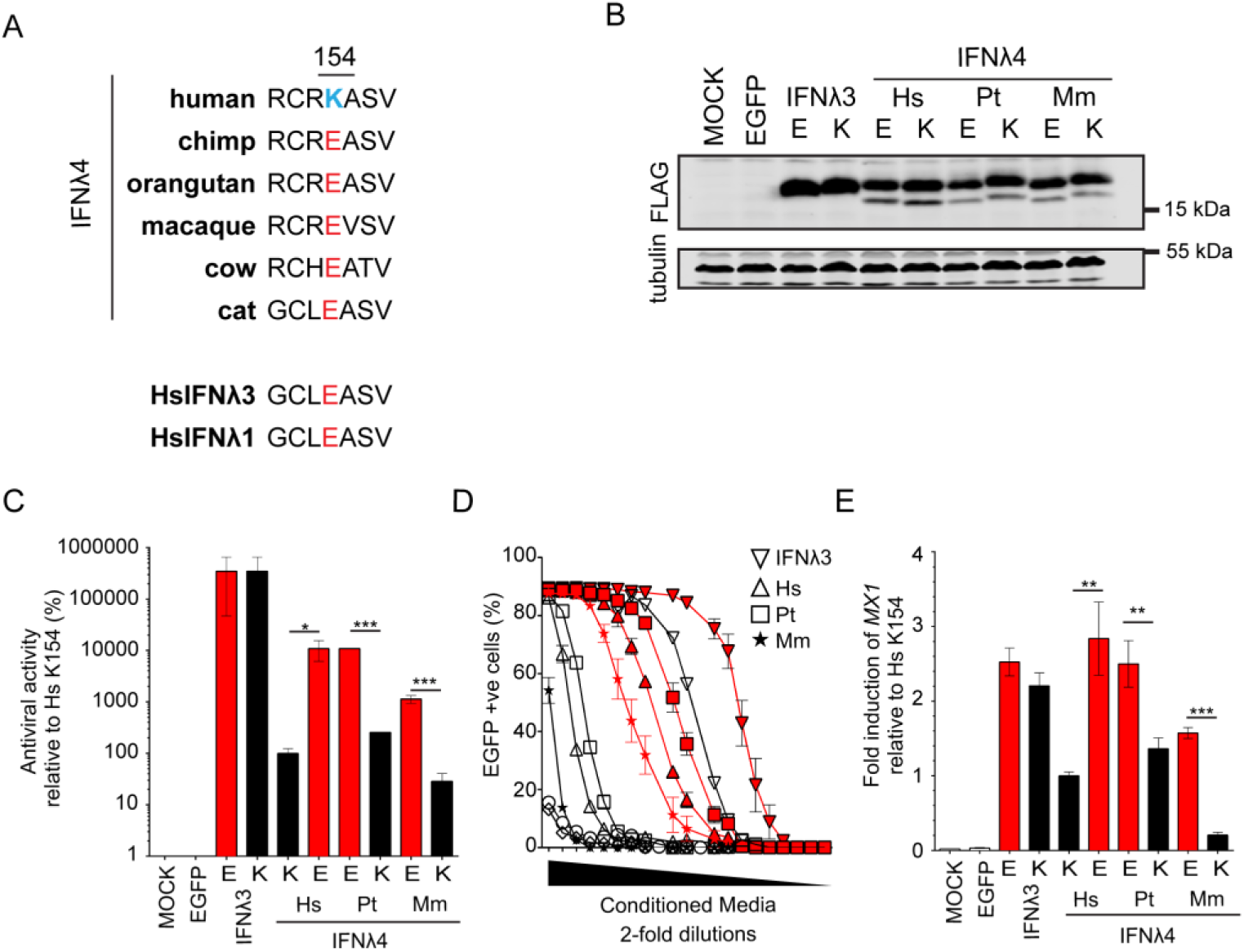
Human IFNλ4 is less active than chimpanzee IFNλ4 due to a substitution at amino acid position 154. Amino acid alignment from positions 151 to 157 for selected orthologues of HsIFNλ4 from different species as well as 2 human paralogues (HsIFNλ1 and HsIFNλ3). At position 154, HsIFNλ4 encodes a lysine (K; blue) while sequences from all other species predict a glutamic acid at this site (E; red). Western blot analysis of intracellular IFNλ4 from different species encoding E or K at position 154 as well as equivalent E and K variants of HsIFNλ3op. HEK293T cells were transfected with the relevant plasmids for 48 hrs prior to preparation of cell lysates. IFNλ4 variants were detected with anti-FLAG antibody (‘FLAG’) and tubulin was used as a loading control. Mock- and EGFP-transfected cells were used as negative controls. (C) EMCV antiviral assay in HepaRG cells of IFNλ from the different species indicated (human [Hs], chimpanzee [Pt] and macaque [Mm]) encoding an E (red bars) or K (black bars) at position 154 alongside the equivalent amino acid substitutions in HsIFNλ3op. Antiviral activity is shown relative to that for HsIFNλ4 in HepaRG cells. Order denotes wt then variant IFNλ for each species. Data show +/- SD (n=3) and are representative of two independent experiments. *** = <0.001; * = <0.05 by unpaired, two-tailed Student’s T test comparing 154E and 154K for each IFN. (D) IFN signalling reporter assay for mutant IFNλ4s from different species encoding an E (red lines) or K (black lines) at position 154 alongside the equivalent amino acid substitutions in HsIFNλ3op. The assay used EGFP-expressing ISG15 promotor HepaRG cells generated by CRISPR-Cas9 genome editing (HepaRG.EGFP-ISG15 cells; clone G8). HsIFNλ4 = triangles; PtIFNλ4 = squares; MmIFNλ4 = stars; HsIFNλ3 = inverted triangles. Serial two-fold dilutions of CM (1:2 to 1:2097152) were incubated with the cells for 24 hrs and EGFP-positive cells (%) were measured by flow cytometry at each dilution. Data shown are average +/- SEM of biological replicates (n=3) and are representative of two independent experiments. Comparison of all E versus K substituted forms of IFNλ4 within a homologue yielded significant values (p = <0.001 by Two-way ANOVA). (E) *MX1* gene expression measured by RT-qPCR for IFNλ4 from different species encoding an E (red bars) or K (black bars) at position 154 alongside the equivalent amino acid substitutions in HsIFNλ3op. Data represent the relative fold change of MX1 mRNA by RT-qPCR in cells stimulated with CM (dilution 1:4) for 24 hrs compared to HsIFNλ4 wt. Data show average +/- SEM (n = 6) combined from two independent experiments. *** = <0.001; ** = <0.01 by unpaired, two-tailed Student’s T test comparing 154E and 154K from each species. Numerical data used for graph construction available in **Supplementary Data File 4 sheet 3.**

Although intracellular expression levels of each IFNλ4 variant were similar, (**Fig 2B**), wt PtIFNλ4 was significantly more active than HsIFNλ4 in each assay and had approximately equivalent activity to the HsIFNλ4 K154E variant in signalling as well as antiviral assays (**Fig 2C-E**). Converting PtIFNλ4 to encode the E154K variant significantly decreased activity to levels that were similar to those for wt HsIFNλ4 (encoding lysine at position 154). Extending the analysis to include rhesus macaque IFNλ4 (*Macaca mulatta*, MmIFNλ4) gave the same pattern whereby wt MmIFNλ4 with E154 had greater activity than its K154 variant. However, wt MmIFNλ4 was less active than either the human or chimpanzee IFNλ4 with E154 indicating that other genetic differences likely modified MmIFNλ4 activity in our assays. Introducing a lysine into HsIFNλ3 had a minimal effect on its activity (**Fig 2C-E**). Overall, we observed a similar ∼100-fold enhancement of activity for E154 over K154 for each of the IFNλ4 orthologues in anti-EMCV activity and EGFP IFN reporter induction. Thus, we conclude that wt HsIFNλ4 has attenuated activity principally because of a single amino acid change at position 154.

### Comparison of the spectrum of antiviral activity of HsIFNλ4 E154 versus K154

To broaden analysis of the impact of a lysine residue compared to a glutamic acid at position 154 in HsIFNλ4, antiviral assays were conducted with other human viruses that are less sensitive to exogenous IFN compared to EMCV, and on different cell lines. Specifically, we used HCV infection in Huh7 cells as well as infectious assays with influenza A virus (IAV) and Zika virus (ZIKV) in A549 cells against single high dilutions of each IFN. As controls, we also included the less active P70S and L79F HsIFNλ4 variants alongside HsIFNλ3op in these assays. Using the HCVcc infectious system in Huh7 cells, HsIFNλ4 K154E significantly decreased both viral RNA abundance compared to wt protein and exhibited a trend towards a lower number of infected viral antigen (NS5A)-positive cells (**Fig 3A**, upper and lower panels respectively). Furthermore, we performed assays examining HCV entry (HCV pseudoparticle system [HCVpp]), viral RNA translation and RNA replication (both assessed with the HCV sub-genomic replicon system). There was no significant difference in the efficiency of HCVpp infection between wt HsIFNλ4 and any of the three variants tested in the MLV-based pseudoparticle assay (**Fig 3B**, upper panel). However, we did observe a greater inhibition when the non-HCV E1E2-containing PPs were used, potentially reflecting the higher efficiency of HCVpp entry compared to non-glycoprotein-containing retroviral PPs that could saturate an inhibitory response (**Fig 3B**, lower panel). wt HsIFNλ4 reduced HCV RNA replication compared to EGFP and introducing the K154E mutation into wt HsIFNλ4 gave a further significant reduction in replication. (**Fig 3C**, upper panel). However, primary translation of input viral RNA was not affected by HsIFNλ addition (**Fig 3C**, lower panel). Thus, HsIFNλ4 reduces HCV RNA replication and the K154E variant exerts greater potency against this stage in the virus life cycle. HsIFNλ4 K154E also reduced titers of IAV and ZIKV to a greater extent than wt protein in A549 cells (∼10-fold; **Fig 3D and E**). Although this was only statistically significant in the context of IAV, a similar trend was evident with ZIKV for the K154E variant compared to wt HsIFNλ4. We found that the P70S and L79F variants consistently reduced the ability of wt HsIFNλ4 to protect against infection in most assays. Taken together, our data further confirmed the greater antiviral activity associated with converting a lysine residue at position 154 in HsIFNλ4 to a glutamic acid residue.

**Fig 3.**
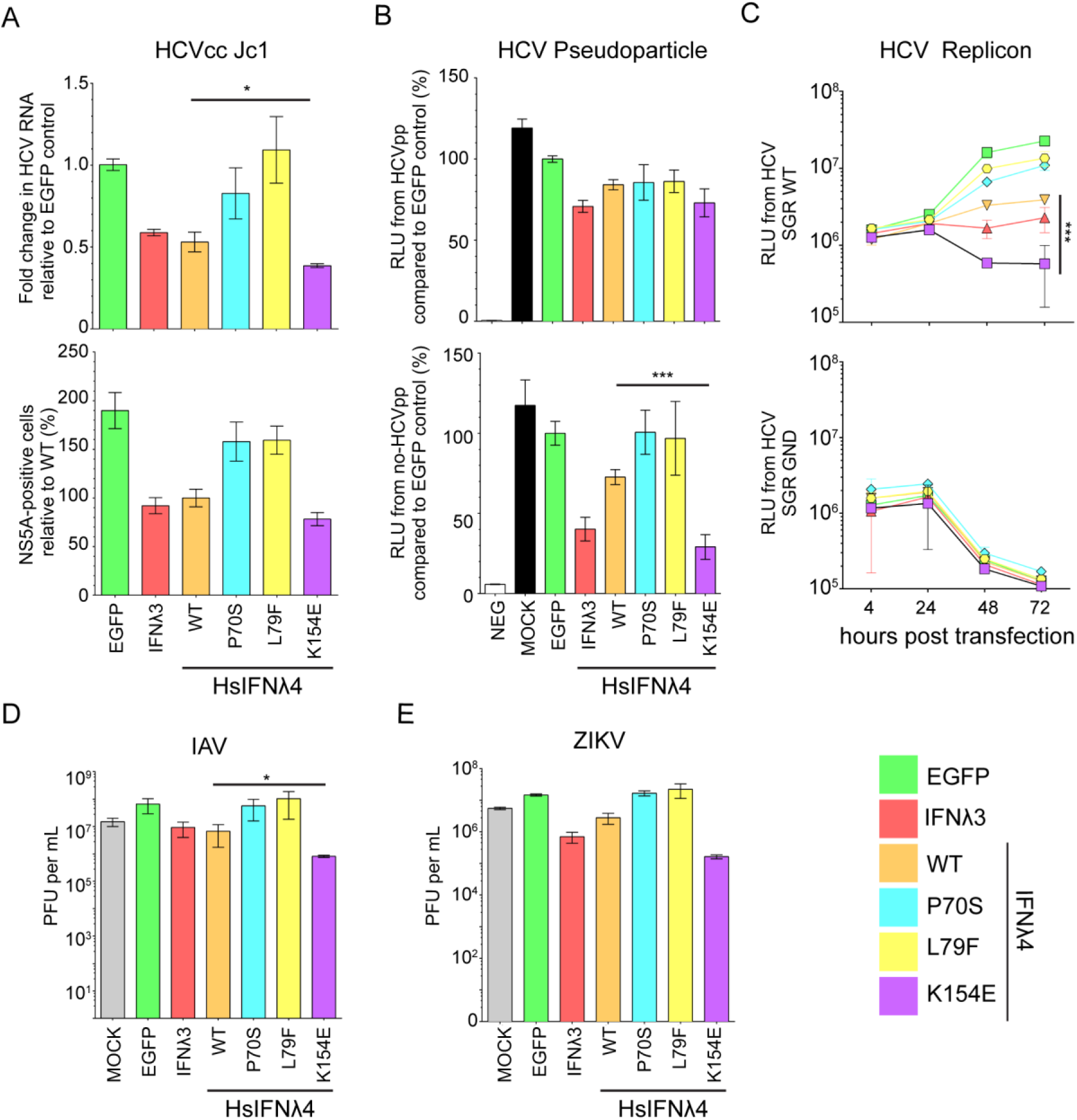
HsIFNλ4 K154E has greater antiviral activity compared to wt HsIFNλ4 K154. (A) Antiviral activity of HsIFNλ4 variants against HCVcc infection in Huh7 cells measured by RT-qPCR of viral RNA (upper panel) and virus antigen-positive cells (HCV NS5A protein; lower panel). HsIFNλ*-*containing CM (1:3) was incubated with Huh7 cells for 24 hrs before infection with HCVcc Jc1 (MOI = 0.01). HCV RNA was measured by RT-qPCR on RNA isolated at 72 hpi. Results shown are relative to infection in cells treated with EGFP CM (upper panel) or wt HsIFNλ4 (lower panel). Data show +/- SEM (n=6) combined from two independent experiments. * = <0.05 by unpaired, two-tailed Student’s T test comparing wt and K154E. (B) The effect of HsIFNλ4 variants on JFH1 HCV pseudoparticle (pp) infectivity in Huh7 cells. Relative light units (RLU) in the lysate of luciferase-expressing MLV pseudoparticles following inoculation of Huh7 cells stimulated with CM (1:3), relative (%) to CM from EGFP-transfected cells. The upper panel shows data from MLV pseudoparticles containing JFH1 glycoproteins E1 and E2 while the lower panel indicates data from MLV pseudoparticles that lack E1 and E2 (MLV core particles). Luciferase activity was measured at 72 hrs after inoculation. Error bars show +/- SEM (n=6). (C) The effect of HsIFNλ4 variants on transient HCV RNA replication (upper panel) and translation (lower panel) using a subgenomic replicon assay in Huh7 cells. Huh7 cells were treated with CM (1:3) for 24 hrs before transfection with *in vitro* transcribed JFH1 HCV-SGR RNA expressing *Gaussia* luciferase; the upper and lower panels show data from wt (replication competent) and GND (non-replicative) sub-genomic replicons respectively. RLU secreted into the media was measured at 4, 24, 48 and 72 hpt. Error bars show +/- SD (n=3). Data are representative of two independent experiments. *** = <0.001 by two-way ANOVA on wt versus K154E. Comparing wt and K154E RLU at 72h by two-tailed Student’s T-test gave a significant difference ** = <0.01. Antiviral activity of wt and variant HsIFNλ4 on IAV (WSN strain) (D) or ZIKV (strain PE243).(E) infection in A549 cells as determined by plaque assay of virus released from infected cells at 48 hpi for IAV or 72 hpi for ZIKV. HsIFNλ4-, HsIFNλ3- and EGFP*-*containing CM (1:3) was incubated with A549 cells for 24 hrs before infection with IAV strain (MOI = 0.01 PFU/cell). Supernatant was harvested and titrated on MDCK cells for IAV or Vero cells for ZIKV. Error bars show +/- SEM (n=3). * = <0.05 by unpaired, two-tailed Student’s T test comparing wt and K154E. Numerical data used for graph construction available in **Supplementary Data File 4 sheet 4.**

### Transcriptomic analysis of cells stimulated with HsIFNλ4 variants

The enhanced antiviral activity of HsIFNλ4 E154 against multiple viruses in different cell lines suggested that this variant may differentially affect global transcription of antiviral ISGs. To test this hypothesis and examine the impact of HsIFNλ4 on global transcription, A549 cells were treated with wt and variant forms of HsIFNλ4 that had different antiviral activities and transcriptional changes were analysed by RNA-Seq at 24 hrs post stimulation (**Fig 4**). A549 cells were used because they recapitulate the functional differences in HsIFNλs as observed in other cell types and are widely used as a cell line model for epithelial antiviral immunity. The data revealed that K154E induced the broadest profile of significantly differentially-regulated genes (n=273) compared with either the wt protein (n=178) or the P70S variant (n=115; **Fig 4A – C and Supplementary Data File 2**). The pattern of genes induced by the positive control HsIFNλ3op and HsIFNλ4 K154E were very similar (**Fig 4B and C)**. From IPA pathway analysis, all HsIFNλs induced the same transcriptional programmes with differences in the overall significance of these pathways, most notably enhancement of the antigen presentation and protein ubiquitination pathways with the K154E variant (**Fig 4D**). Many of the differentially-expressed genes shared by HsIFNλ4 wt, K154E and P70S included known restriction factors with antiviral activity (e.g. *IFI27, MX1, ISG15;* **Fig 4E**) although the magnitude of induction was consistently greatest for HsIFNλ4 K154E (**Fig 4F**). There were also several ISGs that only achieved significant induction by K154E and HsIFNλ3op (e.g. *IDO1*, *IRF1* and *ISG20*; **Fig 4E and F**). We predict that the apparent selectivity by IFNλ4 K154E results from the greater potency of this variant compared to wt HsIFNλ4 and the P70S variant allowing genes to reach the significance threshold (**Fig 4F**). Enhanced production of antiviral genes in cells treated with HsIFNλ4 E154 would explain differences in antiviral activity against EMCV, HCV, IAV and ZIKV.

**Fig 4.**
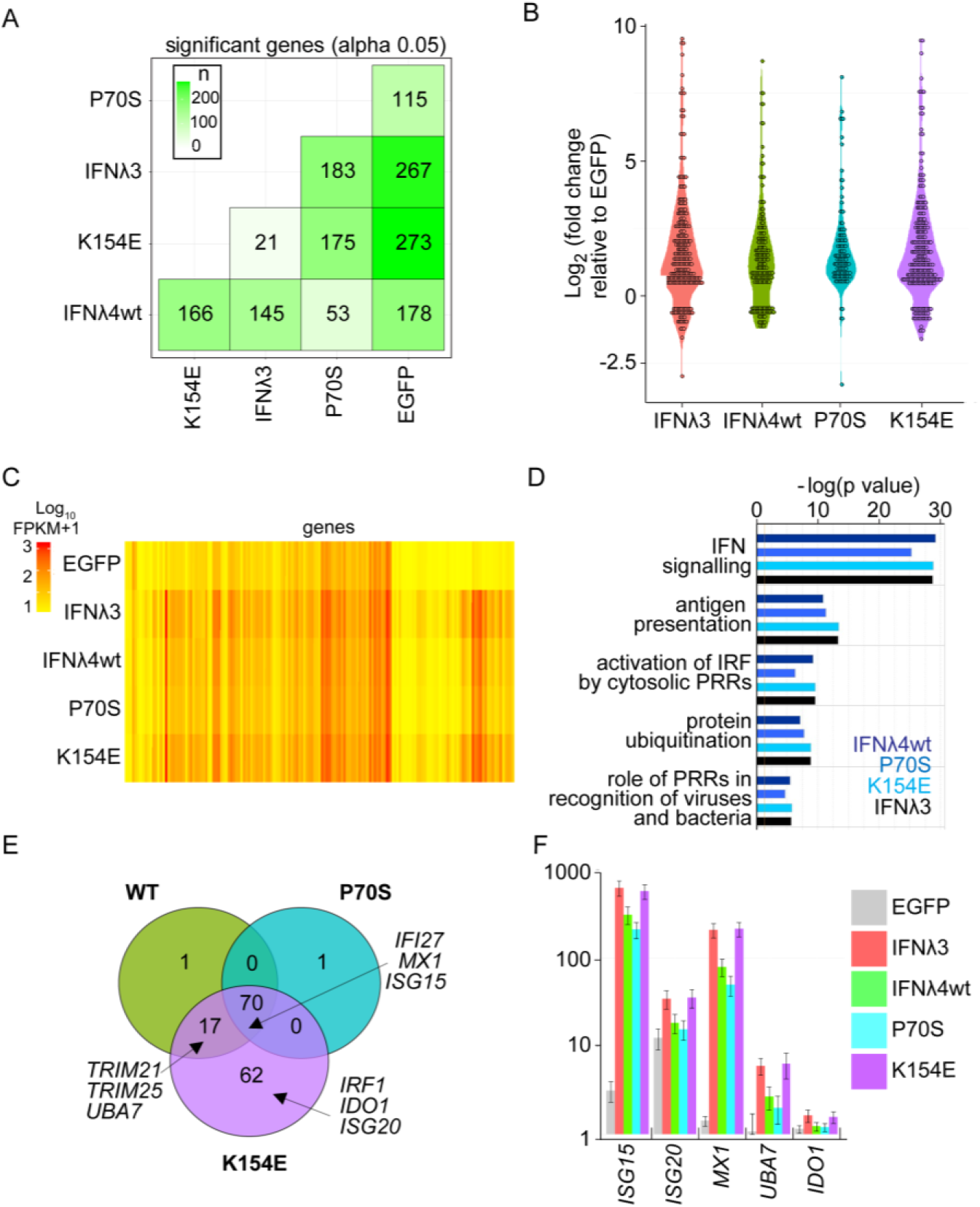
HsIFNλ4 E154 induces more robust antiviral gene expression than the wt K154 variant. A549 cells were treated with CM (1:3 dilution) for 24 hours from cells transfected with plasmids expressing the different HsIFNλs and EGFP. After isolation of RNA, transcriptome analysis was carried out by RNA-Seq. (A) Total number of significantly differentially-expressed genes in each experimental condition (x-axis) relative to each other condition (y-axis). Colour shaded by differences in numbers of transcripts between sample 1 (x-axis) and sample 2 (y-axis) are shown. (B) Violin plot of all significant, differentially-expressed genes (over two-fold) (log2 fold change for each condition compared to RNA from cells treated with EGFP). CM was obtained from cells transfected with HsIFNλ3op (red); HsIFNλ4 wt (green); HsIFNλ4 P70S (cyan), and HsIFNλ4 K154E (purple). (C) Heat map of all significantly differentially-expressed genes (over two-fold) (log10 Fragments Per Kilobase of transcript per Million mapped reads (FKPM) in each experimental condition including EGFP CM-stimulated cells. Genes shown as columns and values are not normalised to negative control. (D) Pathway analysis using IPA on all significantly differentially-expressed genes (>2 fold) for each variant compared to EGFP. The top five most significantly induced pathways are shown [-log(p value)]. (E) Comparison of differentially-expressed genes (significant and at least 2-fold difference) stimulated by the HsIFNλ4 variants (HsIFNλ4 wt in green, HsIFNλ4 P70S in cyan and HsIFNλ4 K154E in purple) illustrated by a Venn diagram showing shared and unique genes. Three examples in overlapping and unique areas of the Venn diagram are highlighted. (F) Raw gene expression values (FKPM+1) for representative genes from core, shared and K154E-‘specific’ groups for the different treatments. Data are shown as mean +/- SD (n=3). Exemplary genes selected were: *ISG15* and *MX1* (core), *UBA7* (not significantly induced by P70S), and *ISG20* and *IDO1* (apparently specific for K154E). All transcriptomic analysis and gene lists are available in **Supplementary Data File 2.**

### Comparison of human and chimpanzee intrahepatic gene expression during HCV infection

Direct *in vivo* validation of our transcriptomic findings alone on the enhanced activity of the K154E variant would require liver biopsy samples from either HCV-infected Pygmies or chimpanzees combined with equivalent samples from infected humans encoding wt HsIFNλ4. This was not possible since such tissue samples are not available from the Pygmy population infected with HCV and biopsies from acutely infected individuals are exceptionally rare. Moreover, chimpanzees are no longer used for experimental studies for ethical reasons. Therefore, we compared lists of reported differentially-expressed genes during acute HCV infection in humans and chimpanzees from the available literature. In the case of humans, there is only one report that analyses the transcriptional response in acute infection (31). For chimpanzees, gene expression analysis is available from four independent studies (32–35) which include longitudinal data from serial biopsies. Therefore, all of the data was collated and we focused our comparisons on periods when human and chimpanzee biopsies were taken across the same time period after initial HCV infection (between 8 and 20 weeks post infection).

Comparative gene expression analysis revealed distinct host responses in humans and chimpanzees as well as overlapping differentially-regulated genes (**Fig 5A and Supplementary Data File 3**). In chimpanzees, the transcriptional profile contained significantly expressed genes that were type I/III IFN-regulated ISGs known to restrict HCV infection (*RSAD2, IFI27* and *IFIT1*) (2), as well as genes involved in antigen presentation and adaptive immunity (*HLA-DMA* and *PSMA6*). These genes were not significantly differentially expressed in humans, whose response was mainly directed towards up-regulation of pro-inflammatory genes (for example*, CXCL10, CCL18* and *CCL5*) and metabolism genes (*AKR1B10* and *HKDC1*) (**Fig 5A and Supplementary Data File 3**). This was consistent with previous characterisation of the human acute response to HCV infection that failed to detect a major type I/III IFN signature but predominantly found a type II or IFN-gamma-mediated response (31). From the available longitudinal data, the ‘chimpanzee-biased’ differentially-expressed genes were induced early in infection and remained significantly up-regulated during the acute phase following an early peak after infection (**S6 Fig A and B**). Differences were also reflected in pathway analysis in terms of the most significant pathways and their overall levels of significance (**Supplementary Data File 3**). For example, the ‘chemokine-mediated signalling pathway’ was upregulated in humans but not chimpanzees whereas the T cell receptor signalling pathway which was modulated in chimpanzees was not significantly altered in humans. Inspection of the raw data from humans indicated that many apparently ‘chimp-biased genes’ were expressed but did not reach significance in the original study. These genes were typically induced at a lower level in the human group when compared to averaged values for chimp studies across the similar time period (**Fig 5B**). Furthermore, there was a greater induction of antiviral ISGs in chimpanzees during chronic infection in comparison to humans although to a less pronounced effect (**S6 Fig C**).

**Fig 5.**
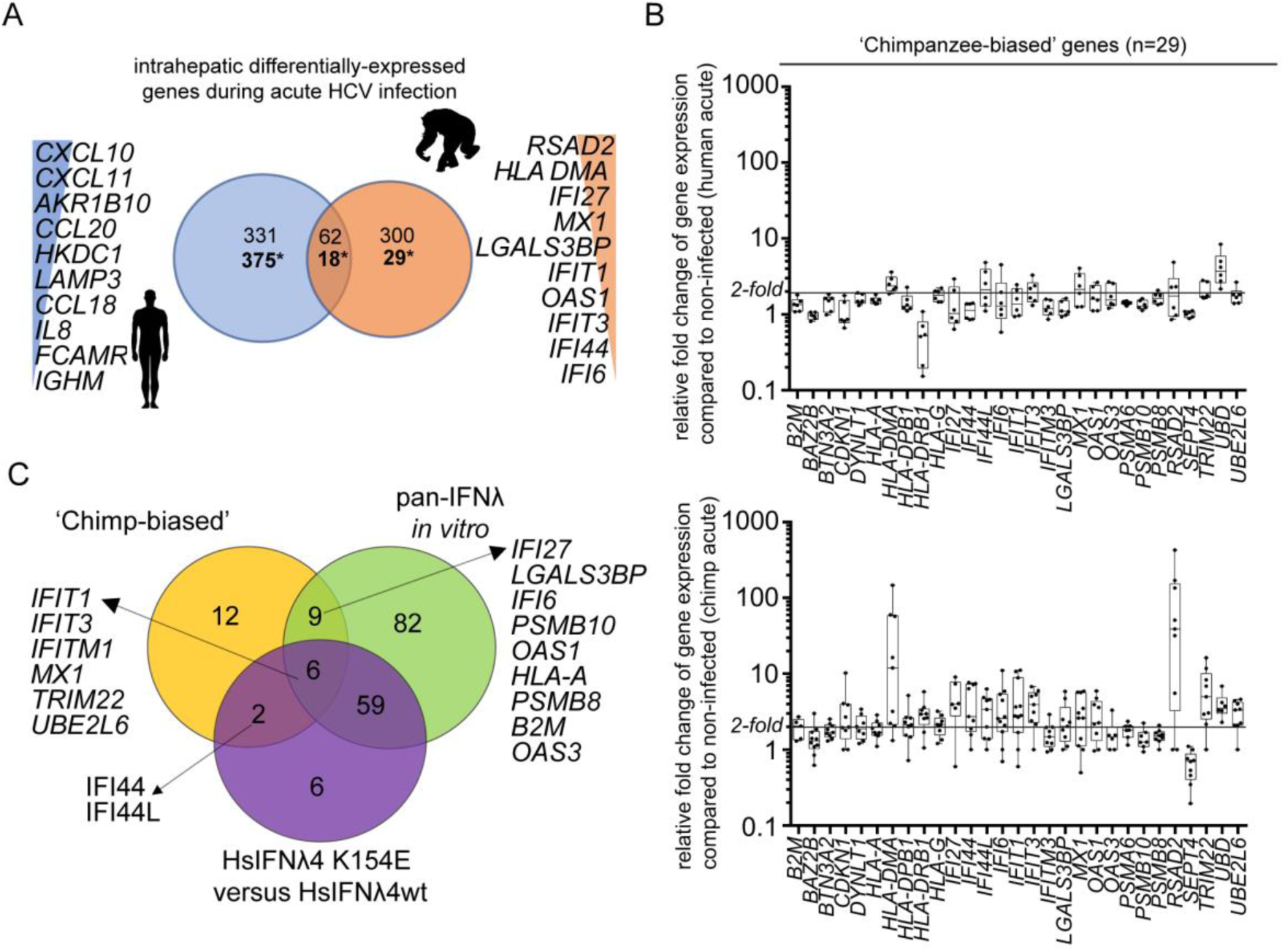
Chimpanzees induce greater levels of antiviral ISG expression during HCV infection *in vivo*. (A) Numbers of shared and unique differentially-expressed genes in liver biopsies from HCV-infected humans (blue) and experimentally-infected chimpanzees (orange) during the acute phase of infection represented as a Venn diagram (also see Supplementary Data File 1). Gene expression during a time period of between 8 and 20 weeks was used where comparable published data for both species exists. The top ten species-‘specific’, differentially-expressed genes are shown ranked by levels of expression. Two sets of values for each comparison are shown; above shows the total differentially-expressed genes from at least one study while * highlights the value relating to the ‘core’ chimpanzee analysis that considered only the genes differentially-expressed in at least two studies of chimpanzee acute HCV infection. (B) Fold change of expression compared to controls (two uninfected individuals) for the 29 ‘chimp-biased’ genes in humans (upper) and chimpanzees (lower) shown as box plot and whiskers. Data are shown as box and whiskers to indicate median and range. Each value is illustrated by a black circle. The chimpanzee values represent an average of all fold changes for each chimpanzee over the time period. (C) Venn diagram analysis comparing the 29 chimpanzee-biased genes to the RNA-Seq data for all IFNs (GFP versus IFN) and for HsIFNλ4 K154E versus wt specifically. Illustrative gene names are shown as examples. All data are available in **Supplementary Data File 2.**

From examining the *in vivo* biopsy data, we identified a group of 29 chimpanzee-biased genes in liver biopsies that were up-regulated during acute infection to a greater extent compared to humans. Comparing this set of genes to those from the RNA-Seq transcriptomic data obtained *in vitro* (**Fig 4)** showed that the majority (17/29 genes) of the chimpanzee-biased genes were induced by HsIFNλ4 stimulation, with approximately half (8 genes) of those being significantly up-regulated to a great extent with K154E compared to wt, including *MX1*, *IFITM1*, *IFIT1*, *IFIT3*, *TRIM22* and *IFI44L* (**Fig 5C**). Thus, there are similarities between our *in vitro* analysis and published *in vivo* studies that would correlate with differences in IFNλ4 activity between humans and chimpanzees.

### Mechanism of action for the enhanced activity of the HsIFNλ4 E154 variant

Having established the greater antiviral potential for the E154 IFNλ4 variant and its apparent evolutionary relevance, we set out to determine the possible basis for its enhanced activity. No crystal structure for HsIFNλ4 is available but a homology model based on comparison with the IFNλ3 structure has been reported (24). We expanded this predicted model based on both of the IFNλ1 and IFNλ3 crystal structures to explore the possible impact of K154E, P70S and L79F on IFNλ4 function (**Fig 6A and S8 Fig**; (36, 37)). As has been previously described, the sequences in helix F, which binds to IFNλR1, are relatively well conserved (18, 24) The position equivalent to amino acid 154 in IFNλ4 is a glutamic acid in both IFNλ1 and IFNλ3 (amino acid position 176 in IFNλ1 and 171 in IFNλ3) and its side chain faces inward towards the opposing IL10R2-binding helices C and D (**Fig 6A**). The free carboxyl group of glutamic acid forms non-covalent intramolecular interactions with two non-linear segments on IFNλ1 and 3 (IFNλ1 residue K64, and in IFNλ3 K67 and T108). In IFNλ4, these E154-interacting positions are not conserved compared to IFNλ1/3 although homologous positions do exist with biochemically similar residues (IFNλ4 R60, and R98 that lies just upstream of the residue homologous to IFNλ3 T108).

**Fig 6.**
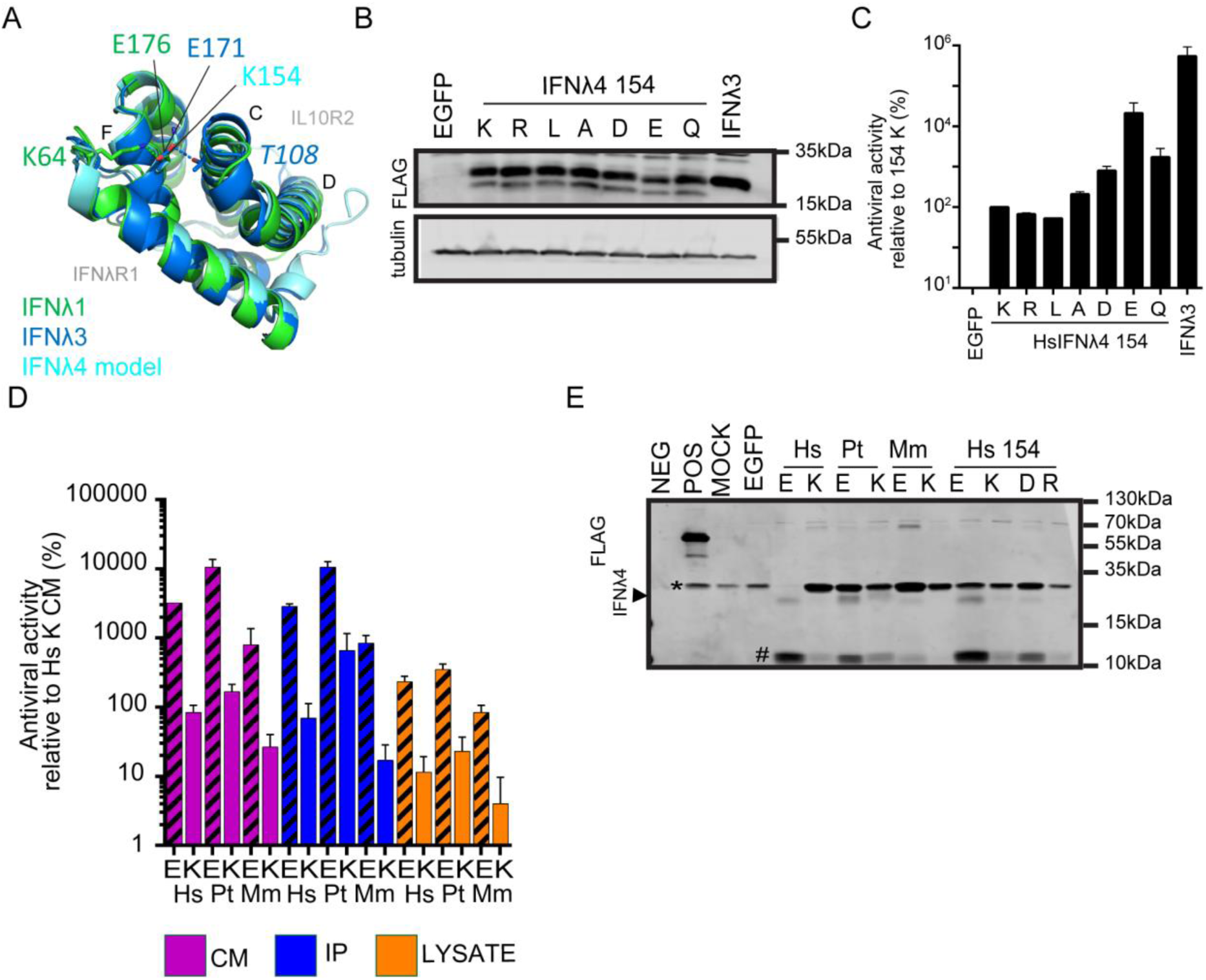
Mechanism of action of the IFNλ4 K154E variant. (A) Modelled structure of HsIFNλ4 showing position 154 at a central location in the molecule with reference to receptor subunit-binding interfaces (IFNλR1 and IL10R2). Overlapping crystal structures for HsIFNλ1 (green) and HsIFNλ3 (dark blue) are overlaid together with a homology model for HsIFNλ4 (light blue). In the overlapping structures, the homologous positions for HsIFNλ4 E154 (E176, IFNλ1; E171, IFNλ3) make intramolecular non-covalent interactions with two distinct regions within IFNλ. (B) Detection of intracellular IFNλ4 154 mutants (K, R, L, A, D, E and Q) by Western blot analysis of lysates from plasmid-transfected producer HEK293T cells. The IFNλ4 variants were detected with an anti-FLAG antibody. Tubulin was used as a loading control. (C) Antiviral activity of HsIFNλ4 IFNλ4 154 mutants (K, R, L, A, D, E and Q) in an anti-EMCV CPE assay relative to CM from wt HsIFNλ4 (K154 variant) in HepaRG cells. Data show mean +/- SEM combined from three independent experiments. (D) Antiviral activity of IFNλ4 found in CM, intracellular lysate and immunoprecipitated CM from the different species indicated (human [Hs], chimpanzee [Pt] and macaque [Mm]) encoding an E or K at position 154 in an anti-EMCV CPE assay relative to CM from wt HsIFNλ4 in HepaRG cells. Data show mean +/- SEM from two independent experiments. (E) Detection of extracellular IFNλ4 from different species as well as select mutants at position 154 (E, K, D and R) by Western blot analysis of samples of FLAG-tag immunoprecipitated CM (1 ml) from plasmid-transfected producer HEK293T cells. A BAP-FLAG fusion protein was used an immunoprecipitation control (POS). The IFNλ4 variants were detected with an anti-FLAG antibody. A FLAG-positive lower molecular weight product, which is potentially a degradation product is highlighted with a #. An upper band running near to the IFN is shown (*) which is likely antibody fragments from the immunoprecipitation reaction. Blot is representative of three independent experiments. Numerical data used for graph construction is available in **Supplementary Data File 4 sheet 5.**

To test whether the biochemical properties of glutamic acid at position 154 contribute to IFNλ4 activity, a panel of variants was constructed with biochemically distinct amino acids (R154, L154, A154, D154 and Q154). Firstly, intracellular expression of each variant at position 154 was approximately equivalent (**Fig 6B**). In signalling assays, the order of activity was E>Q/D>A>L>K>R (**S7 Fig A**). We found a similar pattern in the EMCV antiviral assays except that L154 had the least activity (**Fig 6C**). We interpret these findings to conclude that E154 is biochemically the most favoured residue at this position with regards to antiviral potential, and that substitution of E154 to lysine results in the lowest potency for IFNλ4 activity. Interestingly, both Q154 and D154 had ‘intermediate’ activity compared to E154 and K154, suggesting that side chain length and negative charge are important to maximise the activity of IFNλ4.

In a final series of experiments aimed at giving further insight into the mechanism of action of IFNλ4 K154E, we compared the relative activities and abundance of different IFNλ4 variants in cell lysates (i.e. intracellular protein) and supernatants (i.e. extracellular protein). As wt HsIFNλ4 is poorly secreted into the supernatant from transfected cells in the absence of enrichment (24, 38) IFNλ4 in CM was immunoprecipitated using an anti-FLAG antibody. In antiviral assays, the activity of human, chimpanzee and macaque E154 variants from cell supernatants, IP fractions and lysates was greater than the corresponding K154 variants in agreement with our earlier results (**Fig 6D and Fig 2C-E**). Moreover, the D154 and R154 variants yielded patterns for cell lysates, cell supernatants and immunoprecipitated IFNλ4 protein such that D154 had intermediate activity between E154 and K154 while R154 had approximately equivalent activity to K154 (**S6 Fig B**). Thus, each variant displayed a similar pattern of activity irrespective of the source of IFNλ4. From Western blot analysis, the E154 and K154 variants for each individual species were detected at similar levels in cell lysates (**S6 Fig E**). The HsIFNλ4 D154 and R154 variants were expressed to slightly higher and lower levels respectively compared to E154 and K154 from humans. Paradoxically, we did not find the same pattern in IFNλ4 abundance for immunoprecipitated protein. Thus, we were able to detect greater amounts of the E154 variants for human, chimpanzee and macaque IFNλ4 compared to their K154 variants (**Fig 6E**). It was not possible to reliably detect macaque K154 or human R154 variants. HsIFNλ4 D154 had levels intermediate between the E154 and K154 variants. Quantitatively, the relative abundance of IFNλ4 E154 and K154 variants in cell lysates for any species differed by 1.3 fold yet the approximate fold increase in antiviral activities were significantly greater and on average 16-fold. For the secreted IFNλ4 variants, we found that not only was there a higher abundance of E154 to K154 protein (9-fold), but activity was 41-fold higher for E154 than K154 variants, which results in a significant 3 to 4-fold rise in antiviral activity not explained by protein abundance. With the exception of macaque K154, the FLAG antibody detected a putative breakdown product of about 11kDa in each of the samples, which we presume arose from cleavage by an unknown intracellular protease as it was also detected in cell lysates. The amount of this lower molecular weight product follows the same pattern as the full-length protein in that there is more with E154 than K154 thus cleavage does not explain differences in antiviral activity. Therefore, we conclude that glutamic acid at position 154 promotes greater antiviral potential by enhancing both IFNλ4 secretion from the cell and – to a lesser extent - its intrinsic potency.

## Discussion

In this study we have identified further functional variants of human and non-human IFNλ4 that expand the spectrum of its activity. By comparing IFNλ4 from different species we demonstrate that the genus *Homo* evolved an IFNλ4 gene with attenuated activity (prior to the TT allele), and that the vast majority of extant humans carry an IFNλ4 variant with lower antiviral potential due to a mutation of a single highly-conserved amino acid residue (E154K). Human African hunter-gatherer Pygmies and chimpanzees encode a more active IFNλ4 (E154). We speculate that position 154 in IFNλ4 plays a key role in intramolecular interactions that may facilitate stabilisation of the protein thereby influencing its signalling potential and antiviral activity (**S9 Fig**).

### Implications of the E154K substitution for IFNλ4 evolution

Our analysis suggests that the *Homo* IFNλ4 orthologue acquired the E154K substitution, yielding a less active protein, after the genetic divergence of the hominid *Homo* and *Pan* ancestral lineages (estimated to be at most 6 million years ago in Africa (39)) but before human/Neanderthal divergence (∼370,000 years ago, (40)). Subsequently, the *IFNL4* gene acquired two further variants, the P70S and TT alleles that are now common in the human population (18). Acquisition of each of these alleles either further reduced (P70S) or abolished (TT) IFNλ4 activity. Other rare variants have arisen in humans with little impact on HsIFNλ4 antiviral potential based on our *in vitro* assays, except for variants L79F and K154E, which lower and increase activity respectively. To us, the most intriguing of these variants is K154E, which was found only in rainforest ‘Pygmy’ hunter-gatherers from west central Africa (28). Since this variant was not present in the genetic data for San and Archaic Neanderthal and Denisovan human lineages, we speculate that Pygmy populations likely reacquired K154E following divergence of chimpanzees and humans. However, with the ever-increasing availability of genetic data from ancient and extant human populations, it may be possible to identify other populations carrying the E154 variant.

The factors responsible for divergent functional evolution of the *IFNL4* gene within and between species are not known. It has been demonstrated that loss of *IFNL4* has evolved under positive selection in some human populations thus we speculate that differences in exposure to certain pathogenic microbes has driven evolution of the E154 variant. On the one hand, type III IFN signalling enhances disease and impedes bacterial clearance in mouse models of bacterial pneumonia (41). This suggests that IFNλ4 with a lower activity could be beneficial during non-viral infections although a link between *IFNL4* genotype and bacterial infection in humans has not yet been made. Conversely, we postulate that the presence of more active IFNλ4 exemplified by E154 in Pygmies and chimpanzees may be linked to increased exposure to zoonotic viral infections in the Congo rainforest, such as pathogenic Filovirus infections (42).

### The impact of IFNλ4 functional differences on virus infection

For decades, experimental studies in chimpanzees have provided unique insight into HCV infection (43) but they do not present with identical clinical outcomes as human subjects. For example, chimpanzees have been reported to clear HCV infection more efficiently than humans (44), rarely develop hepatic diseases similar to humans (45), and are refractory to IFNα therapy (46). Moreover, HCV evolves more slowly in infected chimpanzees, possibly due to a stronger immune pressure that reduces replication compared to humans (47). In humans, *IFNL4* genetic variants are associated with, and thought to regulate, each of these characteristics (18,48,49). Although a myriad of factors could explain these phenotypic differences, including differences in antagonism of the immune response by HCV, we propose that the greater antiviral activity of PtIFNλ4 compared to HsIFNλ4 contributes to the distinct responses to HCV infection in the two species.

Acute HCV infection in human cells *in vitro* and chimpanzees *in vivo* selectively stimulates type III over type I IFNs, which are effective at signalling in hepatocytes (50, 51). Notably, there is no apparent type I/III IFN gene expression signature in liver biopsies from humans with acute HCV infection (31). Differences in IFN signalling during HCV infection have been postulated to explain the ability to control HCV infection in cell culture or following IFN-based therapy in humans (52, 53). Our comparative meta-analysis of the available literature revealed enhanced expression of ISGs with anti-HCV activity as well as genes involved in antigen presentation and T cell mediated immunity in chimpanzees compared to humans. Thus, enhanced expression of ISGs in chimpanzee liver due to higher IFNλ4 activity could lead to greater control of viral infection by both inducing antiviral inhibitors and by coordinating a more effective adaptive T cell response, which is critical for clearance and pathogenesis during HCV infection (54).

We would predict that the response to HCV infection in chimpanzees may be similar in Pygmies with the K154E variant. A recent study in Pygmies from Cameroon, including the Baka and Bakola groups, showed low seroprevalence of 0.6% and no evidence of chronic HCV infection (55). Interestingly, infection in non-Pygmy groups in Cameroon has a seroprevalence of ∼17% (56). One explanation for this difference could be higher IFNλ4 activity in populations with the K154E variant, which may enhance HCV clearance.

### How might IFNλ4 E154K reduce antiviral activity?

In our study of three primate orthologues, glutamic acid at position 154 in IFNλ4 provided greater antiviral activity and enhanced its ability to induce antiviral gene expression. A functional comparison of human and chimpanzee IFN□4 orthologues has been explored previously but no significant differences in signalling activity were observed (30). There are substantial differences in the methodologies used in our study and that of Paquin *et al*., which could explain our ability to detect divergent activity, for example size of tag attached to IFNλ4 and dose of protein used in assays.

Our observed functional differences between E154 and K154 did not correlate with levels of intracellular accumulation or glycosylation. However, we did find that the more active E154 variants for human, chimpanzee and macaque IFNλ4 were detected at higher levels in the immunoprecipitated fractions from cell supernatant (CM) compared to the K154 variants; the D154 and R154 variants also were detected at a lower level than E154. Interestingly, wt HsIFNλ4 with K154 is not secreted efficiently compared to wt HsIFNλ3 (24, 38). This was not due to differences in the signal peptide of HsIFNλ4 or HsIFNλ3 but HsIFNλ4 secretion could be ablated if the single N-linked glycosylation site was mutated (24). Our data suggest that position 154 could further regulate IFNλ4 secretion. Aside from the changes in secretion we find evidence that IFN□4s with E154 are more potent than those with K154 when correcting for the difference in amounts of protein. This increase in potency for E154 was detected in both IP protein and lysates. Moreover, we observed that the difference between E and K is greater in the CM or IP fractions than the cell lysate. The reason for this discrepancy could be explained by a number of factors that are outside of the scope of this study. Based on our modelling of the IFNλ4 structure and further mutational analysis, glutamic acid is apparently the optimal residue at position 154.

At the biochemical level, glutamic acid has the capacity to form electrostatic bonds with charged residues in the IFNλ4 protein and moreover it possesses a side chain which could contribute greater flexibility for such interactions. Notably, replacing glutamic acid with either aspartic acid or glutamine gave higher IFNλ4 activity than either non-polar or positively-charged residues. These potential E154-mediated interactions occur in the region of the protein devoid of cysteine-bonds likely making the interaction between helix F (IFNλR1-binding) and the loop connecting helices C and D (IL-10-R2-binding) particularly flexible. The putative greater structural stability facilitated by E154 may inherently increase the structural integrity of IFNλ4 making this variant more competent for secretion and more potent in signalling through the IFNλR1-IL10R2 surface receptor complex. Increased binding to IFN receptor complexes has been shown to enhance signalling by type I IFNs (57, 58). Further biophysical studies using highly-purified recombinant protein measuring affinity and avidity of HsIFNλ4 wt and K154E for each receptor molecule (as in 27, 37) combined with studies on the mechanism of IFNλ4 release will help address these hypotheses.

To conclude, our study further supports a significant and non-redundant role for IFNλ4 in controlling the host response to viral infections yet one whose activity has been repeatedly attenuated during human evolution, commencing with E154K, with the exception of particular African hunter-gatherer groups. Taken together, this provides the foundation for more detailed investigation into the mechanism of action of IFNλ4 and its overall contribution to host immunity in regulating pathogen infection.

## AUTHOR CONTRIBUTIONS

CGGB, EAC and JMcL designed the experiments. CGGB, EAC, ICF, SS and DM conducted the experiments. CGGB, EAC, SS, AdSF, JLM, KCG, SF and ST provided and analyzed data. CGGB and JMcL composed the manuscript. All authors critically reviewed the manuscript.

## ACKNOWLEDGEMENTS

We are grateful to: Chris Boutell, Richard Elliott and Alain Kohl for IAV, EMCV and ZIKV respectively; Takaji Wakita, Ralf Bartenschlager and Arvind Patel for HCV reagents; Sam Wilson and Carol McWilliam Leitch for critically reading the manuscript. SF and ST were supported by NIH grants 1R01DK104339-0 and 1R01GM113657-01. This work was funded by the UK Medical Research Council (MC_UU_12014/1).

## Materials and Methods

### *IFNL* gene sequence analysis

All available human *IFNL4* genetic variation along with associated frequency and ethnicity data for the human population were collected from the 1000 Genomes database available at the time of study (June 2016) (23) (http://browser.1000genomes.org/index.html). The reference sequence for the human genome contains the frameshift ‘TT’ allele and so potential effects of variants on the HsIFNλ4 predicted amino acid sequence were identified manually following correction for the frameshift mutation (TT to ΔG). The effect of all single nucleotide polymorphisms (SNPs) on the open reading frame (ORF) was thus assessed and re-annotated as synonymous or non-synonymous changes resulting in the selection of coding variants reported here. Inspection of whole genome sequence data from African hunter-gatherers was carried out using previously published datasets (28). We remapped the raw reads of six San individuals (four Jul’hoan and two ‡Khomani San) in the Simon Genomic Diversity Project (29) to the human reference genome (hg19) and conducted variant calling using the haplotype caller module in GATK (v3). Two Jul’hoan individuals were heterozygous at rs368234815 (TT/ΔG genotype, **Supplementary Data File 2**). The genotypes of rs368234815 in Neanderthal and Denisovan were extracted from VCF files that were downloaded from http://cdna.eva.mpg.de/denisova/VCF/hg19_1000g/ and http://cdna.eva.mpg.de/neandertal/altai/AltaiNeandertal/VCF/. Neanderthal and Denisovan genetic data contained only ΔG alleles (Supplementary Data File 2). Amino acid sequences for mammalian IFNλ genes were obtained from NCBI following protein BLAST of the wt HsIFNλ4 polypeptide sequence. Multiple alignments of IFNλ amino acid sequences were performed by MUSCLE using MEGA7. Accession numbers of specific IFNλs used in the experimental section of this study were as follows: HsIFNλ1: Q8IU54; HsIFNλ3, Q8IZI9.2; and for IFNλ4: *Homo sapiens* AFQ38559.1; *Pan troglodytes* AFY99109.1; *Macaca mullata* XP_014979310.1; *Pongo abelii* (orangutan) XP_009230852.1, *Bos* taurus (cow) XP_005219183.1, *Felis catus* (cat) XP_011288250.1.

### Structural modelling

The homology model of the HsIFNλ4 structure used in **Fig 6 and S8 Fig** was generated using the RaptorX online server (http://raptorx.uchicago.edu). The resultant HsIFNλ4 structural model was then structurally aligned with both HsIFNλ1 (PDB 3OG6) (36) and HsIFNλ3 (PDB 5T5W) (37). Visualization, structural alignments, and figures were generated in Pymol (The PyMOL Molecular Graphics System, Version 1.8).

### Recombinant DNA manipulation and generation of IFNλ expression plasmids

DNA sequences encoding the ORFs of HsIFNλ4, PtIFNλ4 and MmIFNλ4 (based on accession numbers above) were synthesized commercially with a carboxy-terminal DYKDDDDK/FLAG tag using GeneStrings or Gene Synthesis technology (GeneArt). As a positive control for functional assays, the

HsIFNλ3 ORF was codon optimised (human) to ensure robust expression and antiviral activity, and is termed ‘HsIFNλ3op’. All IFNλ4 coding region sequences were retained as the original nucleotide sequence without optimisation. Synthesized DNA was cloned into the pCI mammalian expression vectors (Promega) using standard molecular biology techniques. At each cloning step, the complete ORF was sequenced to ensure no spurious mutations had occurred during plasmid generation and manipulation. Single amino acid changes were incorporated using standard site-directed mutagenesis protocols (QuickChange site-directed mutagenesis kit [Agilent], or using overlapping oligonucleotides and Phusion PCR).

### Cell lines

A549 (human lung adenocarcinoma), Huh7 (human hepatoma), HEK293T (human embryonic kidney), U2OS (human osteosarcoma), Vero (African Green Monkey kidney) and MDCK (Madin-Darby canine kidney) cells were grown in DMEM growth media supplemented with 10% FBS and 1% penicillin-streptomycin. Non-differentiated human hepatic progenitor HepaRG cells and genome-edited derivatives were cultured in William’s E medium supplemented with 10% of FBS, 1% penicillin-streptomycin, hydrocortisone hemisuccinate (50 µM) and human insulin (4 µg/mL). All cells were grown at 37°C with 5% CO_2_. Cell lines were routinely tested for mycoplasma and no contamination was detected.

### Plasmid transfection and production of functional IFNλ

Plasmid DNA generated from bacterial cultures (GeneJET plasmid midiprep kit, ThermoScientific) was introduced into cells by lipid-based transfection using Lipofectamine 2000 or Lipofectamine 3000 (ThermoFisher) following manufacturer’s instructions. To produce IFN-containing conditioned media (CM) or measure protein production, HEK293T ‘producer’ cells were grown to near-confluency in 12 (∼4 × 10^5^ cells per well) or 6-well (∼1.2 × 10^6^ cells per well) plates and transfected with plasmids (2 µg) in OptiMEM (1-2 ml) overnight. At approximately 16 hours (hrs) post transfection (hpt), OptiMEM was removed and replaced with complete growth media (1-2 ml). CM containing the extracellular IFNλs was harvested at 48 hpt and stored at −20°C before use. Although antiviral activity was observed at 16 hpt, we chose 48 hpt to harvest CM to ensure robust production and secretion of each IFNλ. Intracellular IFNλs also were harvested from transfected cells at 48 hpt. CM was removed and replaced with fresh DMEM 10% FCS (2 ml) and then frozen at −70°C. To prepare cell lysates with IFNλ activity, plates were thawed and the cell monolayer was scraped into the media and clarified by centrifugation (5 minutes [mins] × 300 g) before use. CM or lysates were diluted in the respective growth medium for each cell line before functional testing as described in the text. Two-fold serial dilutions of CM were used in titration of anti-EMCV activity and ability to induce EGFP in an IFN-reporter cell line. Single CM dilutions of 1:4 (HepaRG and A549) or 1:3 (Huh7) were chosen based on initial experiments for gene expression and non-EMCV antiviral activity measurements to allow measurement of both high and low activity variants.

### FLAG-immunoprecipitation

Immunoprecipitation of extracellular FLAG-tagged IFNλ4 present in the supernatant of transfected cells was carried out using an anti-FLAG M2 antibody-bound gel as described by the manufacturer’s guidelines (Sigma Aldrich). Immunoprecipitated IFNs were used in activity assays and for Western blot analysis. Briefly, resin with anti-FLAG antibody (40 µl) and supernatants (1 ml) were thawed on ice. Beads were washed repeatedly in ice cold buffer before being incubated with IFNs in CM for 2 hrs at 4°C while rocking. Bead-bound IFN was pelleted by centrifugation, washed and eluted with FLAG peptide (100 µl). Positive and negative controls were ‘BAP-FLAG’ and buffer only, respectively. Centrifugation conditions were 8,200 × g for 30 sec at 4°C. One quarter (25 µl) of total immunoprecipitated protein was loaded onto gels for Western blot analysis.

### Relative quantification of RNA by reverse transcriptase-quantitative polymerase chain reaction (RT-qPCR)

Total cellular RNA was isolated by column-based guanidine thiocyanate extraction using RNeasy Plus Mini kit (genomic DNA removal ‘plus’ kit, Qiagen) according to the supplier’s protocol. cDNA was synthesised by reverse transcribing RNA (1 µg) using random primers and the AccuScript High Fidelity Reverse Transcriptase kit (Agilent Technologies); the recommended protocol was followed. Relative expression of mRNA was quantified by qPCR (7500 Real-Time PCR System, Applied Biosystems) of amplified cDNA. Probes for *ISG15* (Hs01921425), *Mx1* (Hs00895608) and the control *GAPDH* (402869) were used with TaqMan Fast Universal PCR Master Mix (Applied Biosystems). The results were normalised to *GAPDH* and presented in 2^−ΔΔCt^ values relative to controls as described in the text. HCV genomic RNA was quantified by RT-qPCR as described previously (59).

### Global transcriptomic measurements and pathway analysis

IFN-competent cells (A549) were stimulated with IFN CM (1:4 dilution) in 6-well plates (∼1.2 × 10^6^ cells) for 24 hrs and global gene expression was assessed by RNA-Seq, using three biological replicates per condition. Sample RNA concentration was measured with a Qubit Fluorometer (Life Technologies) and RNA integrity (RIN) was determined using an Agilent 4200 TapeStation. All samples had a RIN value of 9 or above. 1.5 µg of total RNA from each sample was prepared for sequencing using an Illumina TruSeq Stranded mRNA HT kit according to the manufacturer’s instructions. Briefly, polyadenylated RNA molecules were captured, followed by fragmentation. RNA fragments were reverse transcribed and converted to dsDNA, end-repaired, A-tailed, ligated to indexed adaptors and amplified by PCR. Libraries were pooled in equimolar concentrations and sequenced in an Illumina NextSeq 500 sequencer using a high output cartridge, generating approximately 25 million reads per sample, with a read length of 75 bp. 96.3% of the reads had a Q score of 30 or above. Data was de-multiplexed and fastq files were generated on a bio-linux server using bcl2fastq version v2.16. RNA-Seq analysis was performed using the Tuxedo protocol (60). Briefly, reads from 3 replicates per condition were aligned and junctions mapped against the human reference transcriptome hg38 using Tophat2 with the default settings except library type. Transcriptome assembly was performed using Cufflinks supplying annotations from the reference genome hg38 and the differential gene expression was calculated using Cuffdiff. Differential gene expression was considered significant when the observed fold change was ≥2.0 and FDR/q-value was <0.05 between comparisons. Pathway analysis was carried out using Ingenuity Pathway Analysis [IPA] (Ingenuity Systems, Redwood City, CA, USA).

### Western blot analysis

Cell growth media was removed and monolayers were rinsed once with approximately 0.5 ml PBS before lysis using RIPA buffer (ThermoFisher) containing protease inhibitor cocktail (1x Halt Protease inhibitor cocktail, ThermoFisher, or cOmplete™, Mini, EDTA-free Protease Inhibitor Cocktail, Sigma Aldrich) for 10 mins at 4°C before being frozen at −20°C overnight. Lysates were collected into a 1.5 ml sample tube and clarified by centrifugation (12,000 × g for 15 mins). Samples (10 µl) from the soluble fraction were heated to 90°C for 10 mins with 100 mM dithiothreitol (DTT)-containing reducing lane marker at 90°C for 10 mins. Samples were run on home-made 12% SDS-PAGE gels alongside molecular weight markers (Pierce Lane marker, Thermofisher) before wet-transfer to nitrocellulose membrane. Membranes were blocked using a solution of 50% PBS and 50% FBS for 1 hr at room temperature and then incubated overnight at 4°C with primary antibodies in 50% PBS, 50% FBS and 0.1% TWEEN 20. Secondary antibodies were incubated in 50% PBS, 50% FBS and 0.1% TWEEN 20 for 1 hr at room temperature. Membranes were washed four times (5 mins each) following each antibody incubation with PBS containing 0.1% TWEEN 20. After the 4^th^ wash following incubation with the secondary antibody, the membrane was washed once more in PBS (5 mins) and kept in ddH_2_0 until imaging. Primary antibodies to the FLAG tag (1:1000) (rabbit, lot. 064M4757V, LiCor) and α-tubulin (1:10000) (mouse, lot. GR252006-1, LiCor) were used along with infra-red secondary antibodies (LI-COR) to anti-rabbit (donkey [1:10,000], 926-68073) and anti-mouse (donkey [1:10,000], C50422-05) to allow protein visualisation. Pre-stained, Pageruler Plus marker was used to determine molecular weights (ThermoFisher). Membranes were visualised using the LI-COR system on an Odyssey CLX and the relative expression level of proteins determined using LI-COR software (Image Studio).

### Generation and use of IFN reporter cell lines

An IFN reporter HepaRG cell line was generated to measure IFN activity by introducing the EGFP ORF fused to the ISG15 ORF separated by ribosome skipping sites by CRISPR-Cas9 genome editing. We chose to introduce EGFP in-frame to the N-terminus of the *ISG15* ORF since it is a robustly-induced ISG. We also introduced the blasticidin resistance gene (BSD) for selection purposes. BSD, EGFP and ISG15 were separated using ribosome skipping 2A sequences (P2A and T2A). Transgene DNA was flanked by homology arms with reference to the predicted target site. Homology donor plasmids for CRISPR-Cas9 knock-in were generated through a series of overlapping PCR amplifications using Phusion DNA polymerase followed by sub-cloning into pJET plasmid. Plasmids for CRISPR-Cas9 genome editing (wt SpCas9) were generated using established protocols (61) in order to create plasmids that would direct genome editing at the 5’ terminus of the HsISG15 ORF (exon 2). pSpCas9(BB)-2A-Puro (PX459) V2.0 was a gift from Feng Zhang (Addgene plasmid # 62988). All sequences are available by request. HepaRG cells grown in 6 well dishes were co-transfected with CRISPR-Cas9 editing plasmids targeting the beginning of the ISG15 ORF in exon 2 (exon 1 contains only the ATG of the ORF), and homology donor plasmids described above (1 µg each) using Lipofectamine 2000 and the protocol described above. Transfected cells were selected using puromycin (Life Technologies) (1 µg/ml) and blastocidin (Invivogen) (10 µg/ml) until non-transfected cells were no longer viable. Selected cells were cloned by single cell dilution, expanded and tested for EGFP induction following IFN stimulation. Positioning of the introduced transgene was assessed by PCR amplification on isolated genomic DNA from individual clones (data not shown). Primers were designed to include one primer internal to the transgene and another external to the transgene and found in the target loci (sequences available on request). For use as an effective IFN reporter cell line, cells had to demonstrate robust induction of EGFP expression following stimulation with IFN and evidence of specific introduction of the transgene. This study uses clone ‘G8’ of HepaRG.EGFP-BSD-ISG15 cells. We have not tested whether there is a single transgene integration site or multiple ones nor confirmed that the EGFP produced following stimulation by IFNs results from the expression of the specifically-introduced transgene rather than off-target integration, which is theoretically possible. We do not predict this would affect the cells’ ability to act as a reporter cell line. For use in IFN reporter assays, stimulated cells (in 96 well plates stimulated for 24 hrs; ∼5 × 10^4^ cells per well) were washed, trypsinised and fixed in formalin (1% in PBS) at room temperature for 10 mins in the dark before being transferred to a round-bottomed plate and stored at 4°C in the dark until measurement of EGFP fluorescence. Non-stimulated cells were used as negative controls and the change in % EGFP-positive cells was assessed by flow cytometry using a Guava easyCyte HT (Merck Millipore). For fluorescence microscopy, EGFP induction was measured by indirect immunofluorescence of stimulated cells that were fixed and permeabilised on coverslips prior to antibody binding. An EGFP primary antibody (1:1000, rabbit ab290 Abcam) was used followed by a fluorescent anti-secondary antibody (1:500, Goat anti Rabbit Alexa-Fluor, Thermo Fisher, 568nm). Samples were counter-stained using DAPI and visualised with a confocal laser-scanning microscope (Zeiss LSM 710) under identical conditions.

### Production of virus stocks for antiviral assays

Antiviral activity of IFNλs was determined using encephalomyocarditis virus (EMCV), influenza A virus (IAV; A/WSN/1933(H1N1)), Zika virus (ZIKV; Brazilian strain PE243)(62) and HCV (HCVcc chimeric clone Jc1) (63). EMCV was grown on Vero cells followed by titration on U2OS cells by plaque assay. IAV stocks were generated on MDCK cells and titrated by plaque assay on MDCK cells with protease (TPCK-treated trypsin, Sigma Aldrich). ZIKV was titrated on Vero cells by plaque assay. For all plaque assays, cells were grown in 12 or 6-well plates to ∼90% confluency before inoculation with serial 10-fold dilutions of virus stocks in serum-free Optimem. Inoculum remained on the cells for 2 hrs before removal and the monolayers were rinsed with PBS (1x) and semi-solid Avicell overlay (Sigma Aldrich) was added. For EMCV and IAV, 1.2% Avicell was used, diluted in 1x DMEM 10% FCS, 1% penicillin-streptomycin. For IAV titration, TPCK-treated trypsin was added (1 µg/ml). For ZIKV plaque assay, 2x MEM was used instead of 1x DMEM. HCVcc Jc1 was generated as described previously by electroporation of *in vitro* transcribed viral RNA into Huh7 cells and harvested at 72 hrs post electroporation. After filtration of the supernatant, HCVcc Jc1 stocks were titrated by TCID_50_ on Huh7 cells and stored at 4°C before use. HCVcc Jc1 TCID_50_ assays were performed using anti-NS5A antibody (64). Infected cells at 72 hrs post infection were fixed and permeabilised with ice-cold methanol. Cells were rinsed in PBS, blocked with 3% FCS in PBS at room temperature and incubated overnight with mouse monoclonal anti-NS5A antibody (9E10) at 4°C. After removal of the antisera, cells were rinsed 3 times with PBS containing 0.1% TWEEN 20, and then incubated in the dark at room temperature for 1 hr with secondary antibody [Alexa-fluor 488nm anti-mouse (donkey)]. Cells were finally washed with PBS containing 0.1% TWEEN 20 and NS5A-expressing cells were visualized with a fluorescent microscope.

### Antiviral assays

Cells stimulated with IFNλs were infected with viruses at the following multiplicities of infection (MOI): EMCV (MOI = 0.3; added directly to the media); IAV (MOI = 0.01); ZIKV (MOI = 0.01); HCVcc (MOI = 0.05). For IAV, ZIKV and HCVcc, the inoculum was incubated with cells for at 2 (IAV/ZIKV) or 3 hrs (HCVcc) in 0.5–1.0 ml serum-free Opti-MEM/DMEM at 37°C before removal. Cells were rinsed with PBS and then incubated with fresh growth media for the allotted time (24 hrs for EMCV, 48 hrs for IAV and 72 hrs for ZIKV and HCVcc). At the times stated for individual experiments, infected-cell supernatants were harvested and infectivity was titrated by plaque assay. IAV, ZIKV and HCVcc antiviral assays were all carried out in 12 well plates except for measurement of HCVcc infectivity by indirect immunofluorescence, which was measured in a 96 well plate. In the case of EMCV, a cytopathic effect (CPE) protection assay was employed to assess infectivity (26). Here, HepaRG cells were plated in a 96-well plates (∼5 × 10^4^ cells per well) and, when confluent, were incubated with two-fold serial dilutions of CM or lysate for 24 hrs before the addition of EMCV. At 24 hrs post infection with EMCV, media was removed, cell monolayers were rinsed in PBS and stained using crystal violet (1% in 20% ethanol in H_2_0) for 10 mins. Crystal violet stain was then removed and stained plates were washed in water. The dilution of ∼50% inhibition of EMCV-induced CPE was marked visually and the difference determined relative to wt HsIFNλ4.

Luciferase-expressing MLV pseudoparticles containing the E1 and E2 glycoproteins from JFH1 HCV strain were generated as described (65) along with their corresponding JFH1 E1-E2 deficient controls (particles generated only with MLV core) and used to challenge IFNλ-stimulated Huh7 cells. Huh7 cells grown in 96-well plates overnight (seeded at 4 × 10^3^ cells per well) were stimulated with IFNλs for 24 hrs and transduced with HCVpp. 72 hrs later, cell lysates were harvested and luciferase activity was measured (Luciferase assay system, Promega) on a plate reading luminometer.

For HCV RNA replication assays, RNA was transcribed *in vitro* from a sub-genomic replicon (HCV-SGR) expressing GLuc (wild-type and non-replicating GND) (66). *In vitro* transcribed RNA (200 ng) was transfected using PEI (1:1) into monolayers of Huh7 cells in 96-well plates overnight (seeded at 4 × 10^3^ cells per well) that had been stimulated with IFNλs (24 hrs). At the specified time points, total supernatants (containing the secreted GLuc) from treated Huh7 cells were collected and replaced with fresh growth media. 20µl (∼10% of total volume) was used to measure luciferase activity and mixed with GLuc substrate (1x) (50 µl) and luminescence (as relative light units, RLUs) was determined using a luminometer (Promega GloMax). Pierce *Gaussia* Luciferase Flash Assay Kit (ThermoFisher) was used and the manufacturer’s instructions were followed.

### Comparison of human and chimpanzee intrahepatic gene expression during acute HCV infection

Previously published datasets of intrahepatic differentially-expressed genes from liver biopsies were used to compare human and chimpanzee transcriptomic responses to early HCV infection. At first, we used reported lists of differentially-expressed genes between humans and chimpanzees but further validated observations with raw data from human studies. Studies focusing on acute HCV infection (0 to 26 weeks) in humans and chimpanzees were acquired through manual literature search using Pubmed and gene lists were compiled. For chimpanzees, data was acquired from 4 studies (32–35) and one report was employed for human data (31). The study by Dill et al. comprised single biopsy samples from each of six individuals, while *in toto* the chimpanzee studies combined data from ten animals with multiple, serial biopsies. All studies were carried out using similar Affymetrix microarray platforms except Nanda *et al*. who used IMAGE clone deposited arrays. Although similar microarrays measured different numbers of genes we focused on ‘core’ shared genes from chimpanzee studies. Humans were infected with HCV genotype (gt)1 (n=2), gt3 (n=3) and gt4 (n=1) while chimpanzees were experimentally infected with HCV gt1a (n=6), gt1b (n=3) and gt2a (n=1). The human dataset included individuals with IL28B rs12979860 genotypes T/T, C/T and C/C but no association between IL28B genotype and gene expression was noted (31). Gene names and fold-changes were manually converted to a single format (fold change rather than log2 fold change for example) to allow comparative analysis. Human biopsies were taken between two and five months after presumed infection following known needle-stick exposure, and serial chimpanzee biopsies were taken at different time points from between one week and one year after HCV infection. For comparative purposes, differentially-expressed genes in chimpanzees were included if they were detected during a time period overlapping with the human data. We identified a ‘core’ set of chimpanzee differentially-expressed genes (independently characterized in at least two studies) and compared them to the single human transcriptome study data at equivalent time points (between 8 and 20 weeks post-infection). This approach generated a set of core chimpanzee genes (genes found differentially-expressed in at least 2 studies, >2 fold change compared to controls and during the time frame compared to humans) for comparison with the human data. This is reflected in the ten-fold higher numbers of differentially-regulated genes found in the one human study compared to the ‘core’ (reduced) set assembled from four chimpanzee studies. To validate these findings, we used three studies of chronic infection for which data were available (53,67,68), two from humans employing RNA-Seq and one from chimpanzees using microarray measurement. Gene lists were extracted and a core human list was produced and compared to that from chimpanzees. For shared genes, the fold change values were compared for humans and chimpanzees. The ratio of chimpanzee induction to human induction was calculated.

These gene sets were compared to determine their degree of species-specificity or species-similarity using Venn diagram analysis (http://bioinfogp.cnb.csic.es/tools/venny/). The gene lists for humans and core genes for chimpanzees are shown in the Supplementary Data File 1. For the chimpanzee-biased genes, mean expression values were determined at each time point from individual animals.

### Statistical analysis

For non-transcriptomic analysis (transcriptomic analysis is outlined above), Graphpad Prism was used for statistical testing, which included Students’ T test and ANOVA and post-hoc tests (Dunnett’s test) where appropriate as described in figure legends. ****, p=<0.0001; ***, p=<0.001; **, p=<0.01; *, p=<0.05, are used throughout to denote statistical significance.

---

**S1 Fig.**
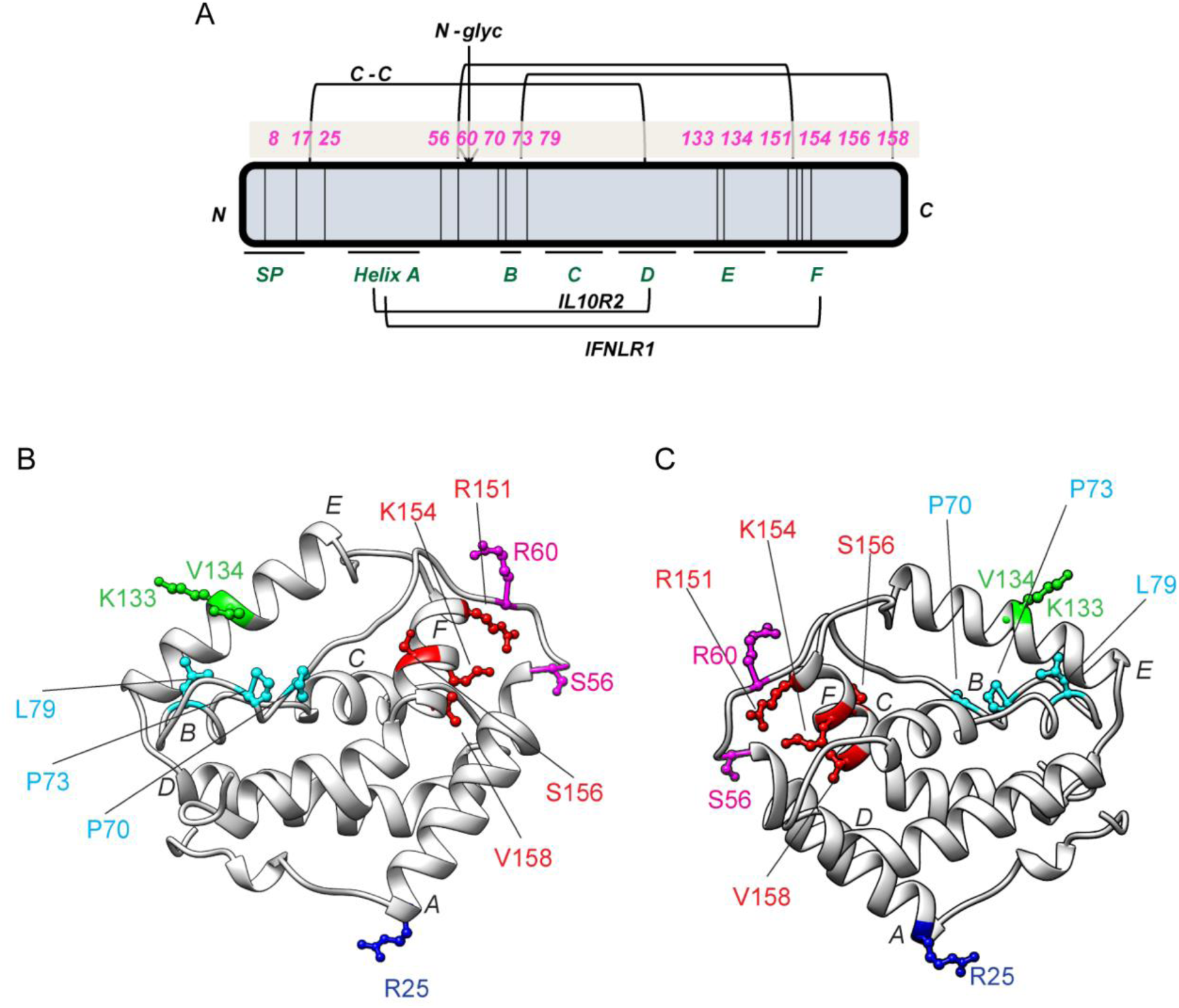
Non-synonymous variants of HsIFNλ4 are located in regions of functional significance. (A) Schematic location of non-synonymous variants in the HsIFNλ4 polypeptide (N- to C-terminus) (above schematic in pink). Regions of predicted structural significance are underlined, including the signal peptide (SP), single N-linked glycosylation site (N-glyc, arrowed), helices (A to F) and disulphide bonds (C-C). Note that there are 2 non-synonymous changes at C17 (C17R and C17Y). Helices involved in receptor interactions (IL10R2 and IFNλR1) are highlighted. (B and C) Location of non-synonymous variants on a homology model of HsIFNλ4 (side chains in colour) from two perspectives. Model was generated using the SWISS-MODEL online software. Helices are labelled A to F. Positions are coloured based on spatial clustering in the primary amino acid sequence.

**S2 Fig.**
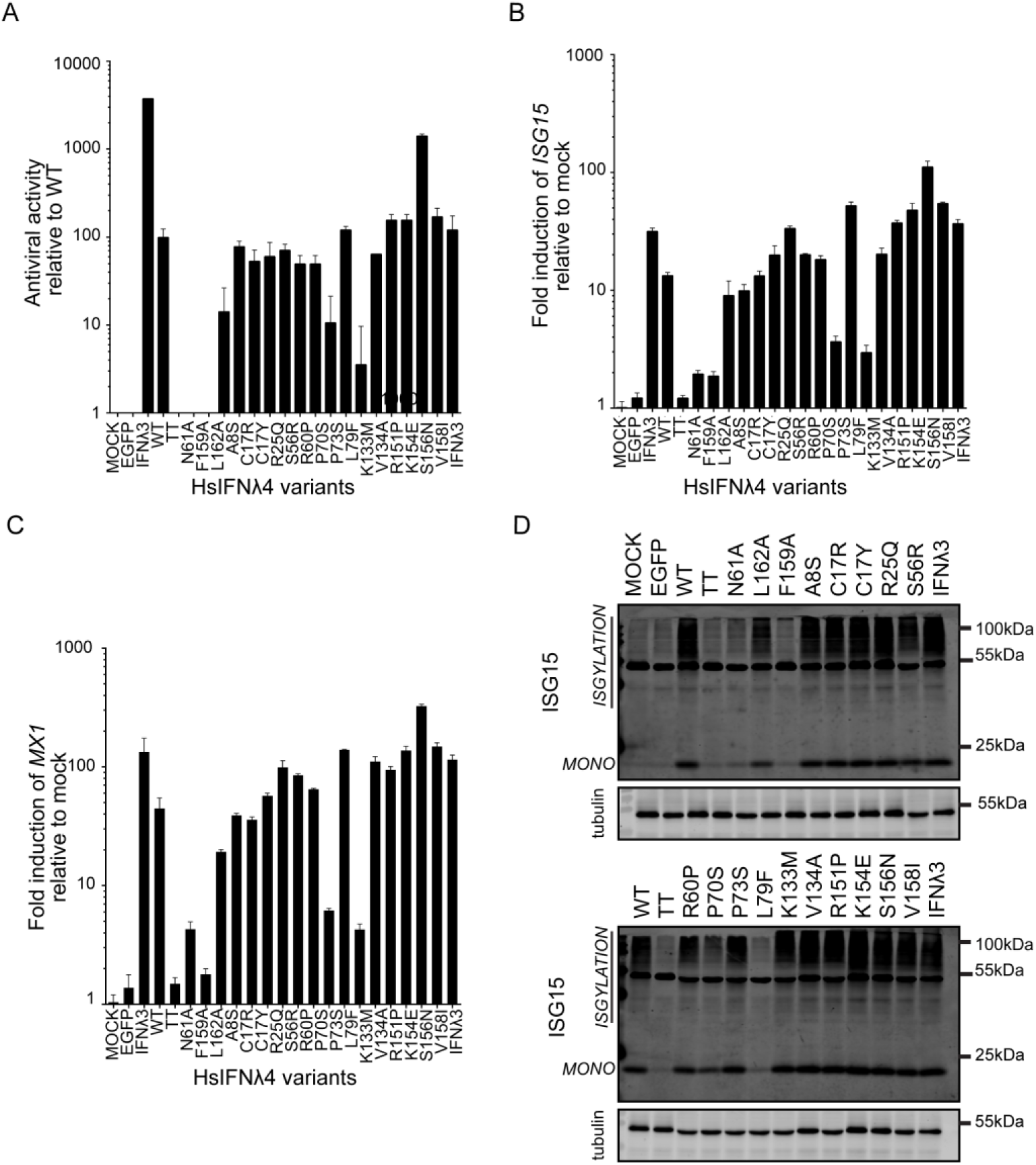
Rare non-synonymous variants of HsIFNλ4 affect antiviral activity. For data shown in panels A-D, all naturally-occurring variants of HsIFNλ4 were tested in antiviral and ISG induction assays. Experimental conditions included a series of controls including HsIFNλ3op (positive control), EGFP and the HsIFNλ4 TT variant (negative controls) as well as non-natural variants of HsIFNλ4 (N61A, F159A, L162A). N61A abrogates glycosylation of HsIFNλ4 while F159A and L162A are predicted to reduce interaction with the IFNλR1 receptor subunit and hence lower activity based on previous studies (27). Panels show data from the following assays: (A) Antiviral activity in an anti-EMCV CPE assay in HepaRG cells. Cells were stimulated with serial dilutions of HsIFNλ4-containing CM for 24 hrs and then infected with EMCV (MOI = 0.3 PFU/cell) for 24 hrs at which point CPE was assessed by crystal violet staining. After staining, the dilution providing ∼50% protection was determined. Data are shown as mean +/- SD of three independent experiments. (B and C) ISG gene expression determined by RT-qPCR following stimulation of cells with HsIFNλ4 variants. Relative fold change of *ISG15* mRNA (B) or *Mx1* (C) in HepaRG cells stimulated with CM (1:4 dilution) from plasmid-transfected cells compared to wt HsIFNλ4. Cells were stimulated for 24 hrs. Error bar represent mean +/- SD (n=3). (D) Western blot analysis of unconjugated and high molecular weight conjugated-forms of ISG15 (‘ISGylation’) from lysates harvested from HepaRG cells stimulated with CM (1:4) for 24 hrs. Numerical data used for graph construction available in **Supplementary Data File 4 sheet 1.**

**S3 Fig.**
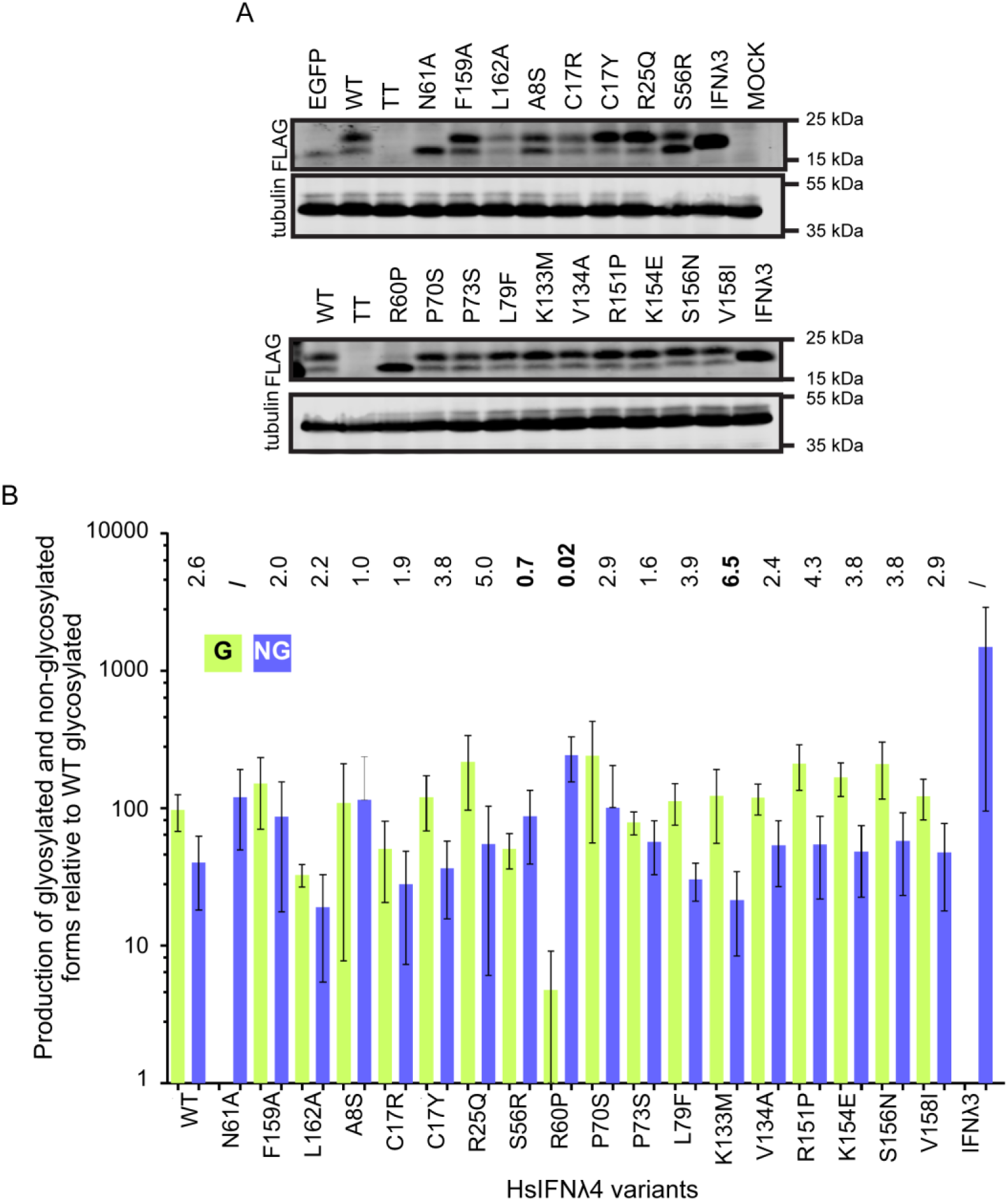
Relative expression of glycosylated and non-glycosylated forms of HsIFNλ4 variants. For data in panels A and B, expression and glycosylation of all naturally-occurring variants of HsIFNλ4 were examined. Experiments included a series of controls including HsIFNλ3op (contains no glycosylation sites), EGFP and the HsIFNλ4 TT variant (negative controls) as well as non-natural variants of HsIFNλ4 (N61A, F159A, L162A). N61A is predicted to abrogate glycosylation of HsIFNλ4. Panel A shows a representative Western blot for the production and glycosylation of HsIFNλ4 variants of lysates from plasmid-transfected producer HEK293T cells as detected with an anti-FLAG (‘FLAG’) primary antibody. Tubulin was used as a loading control. A non-specific band in the EGFP-transfected extract is shown (*). Panel B shows the quantification of intracellular glycosylated (green) and non-glycosylated (blue) HsIFNλ4 variants by Western blot analysis of lysates from plasmid-transfected producer HEK293T cells. Ratio of glycosylated to non-glycosylated is shown above the graph. Two-fold differences from wild-type are highlighted in bold. Data shown are mean +/- SEM combined from three independent experiments. Numerical data used for graph construction available in Supplementary Data File 4 sheet 2.

**S4 Fig.**
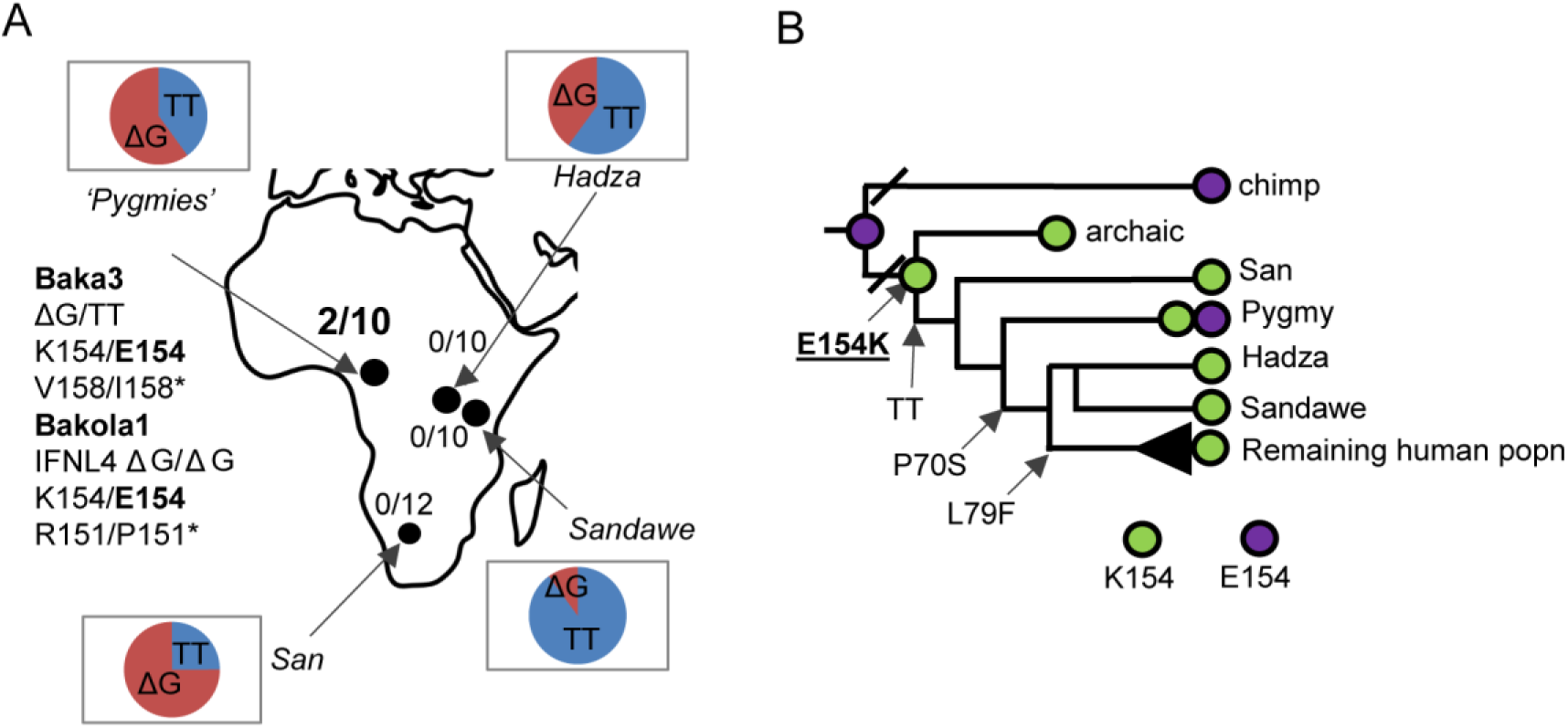
Presence of HsIFNλ4 K154E variant in Pygmies and evolution of HsIFNλ4 variants in human populations. (A) Geographical location and frequency of HsIFNλ4 K154E in African hunter-gatherer alleles (Pygmy, n = 5 individuals, Sandawe (S) n = 5 individuals and Hadza (H) n = 5 individuals). Two Pygmy individuals within two tribes (Baka and Bakola) were found to encode the HsIFNλ4 K154E variant. The proportion of ΔG (red) and TT (blue) *IFNL4* alleles are also shown in pie-charts. (B) Presence of HsIFNλ4 E154 (purple) versus HsIFNλ4 K154 (green) on a cladogram of human and chimpanzee evolution. Archaic human (Neanderthal and Denisovan) as well as other basal human populations (San, Sandawe and Hadza) only encode HsIFNλ4 K154. Earliest detection of the HsIFNλ4 TT frameshift and activity-reducing HsIFNλ4 P70S and HsIFNλ4 L79F variants are shown. All analysis can be found in **Supplementary Data File 1.**

**S5 Fig.**
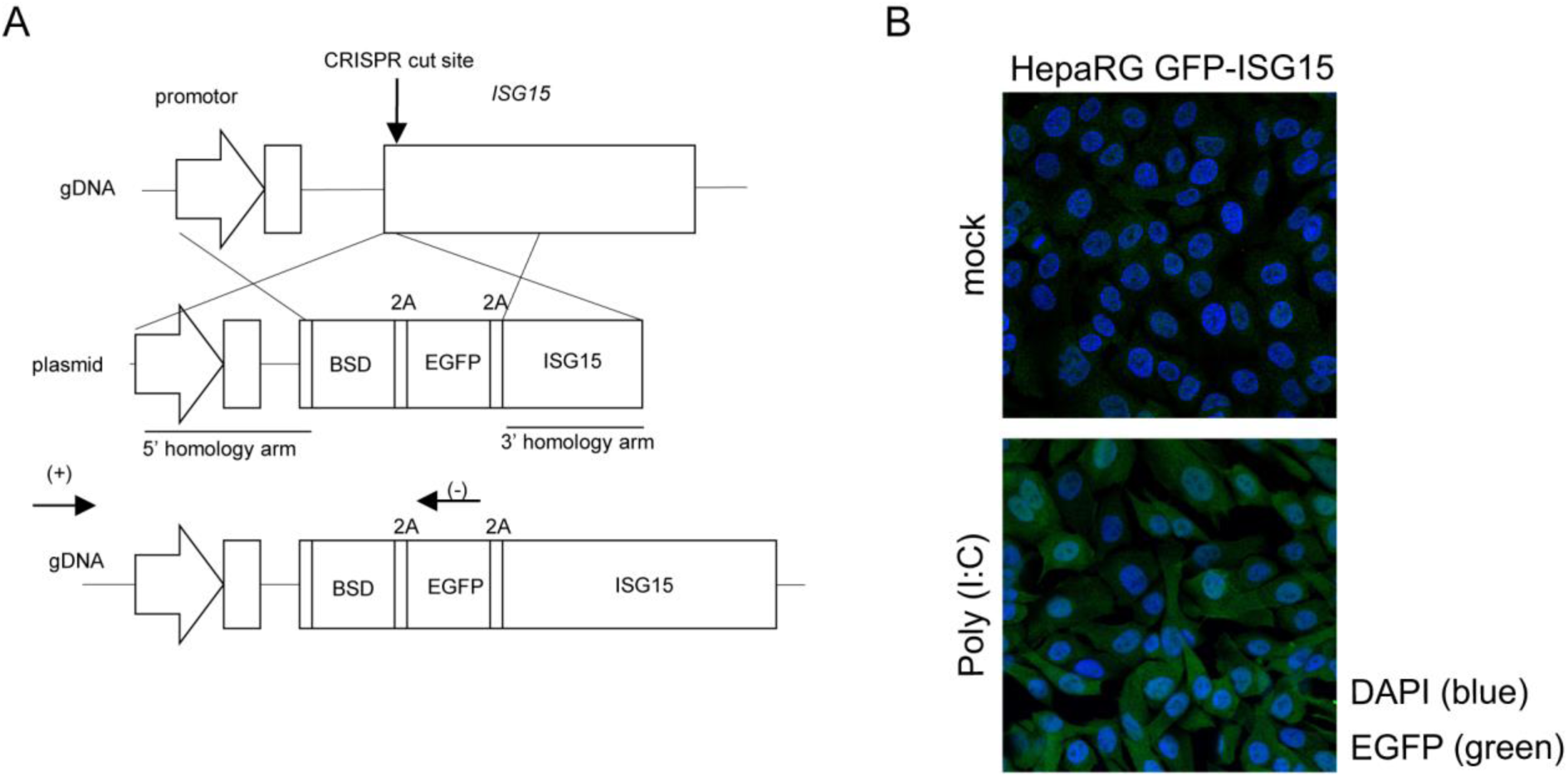
Generation of a reporter HepaRG cell line expressing EGFP in the ISG15 promoter region. (A) Strategy for CRISPR-Cas9 genome editing combined with homologous recombination insertion of DNA sequences to produce an EGFP-expressing ISG15 promotor cell line. The strategy enables the insertion of a cassette in-frame with the ISG15 ORF that encodes blasticidin resistance (BSD) and EGFP genes followed by *ISG15*, separated by ‘2A’ ribosomal skipping sequences. Target gDNA and CRISPR-Cas9 cut sites are shown on the upper cartoon. Binding sites of primers for genotyping are highlighted (+) and (-) in the resulting modified gDNA. Cell lines used for reporters were validated by PCR analysis (data not shown). (B) Induction of EGFP expression (green immunofluorescence) in the G8 cell clone with or without treatment with PolyI:C (1.0 µg/ml for 24 hrs). Poly(I:C) stimulates the Toll-like receptor TLR3, which induces ISG15 expression. Cells were fixed in formalin, permeabilised and stained for indirect immunofluorescence with an anti-EGFP primary antibody before addition of a secondary antibody. DAPI was used as a counter stain for cell nuclei.

**S6 Fig.**
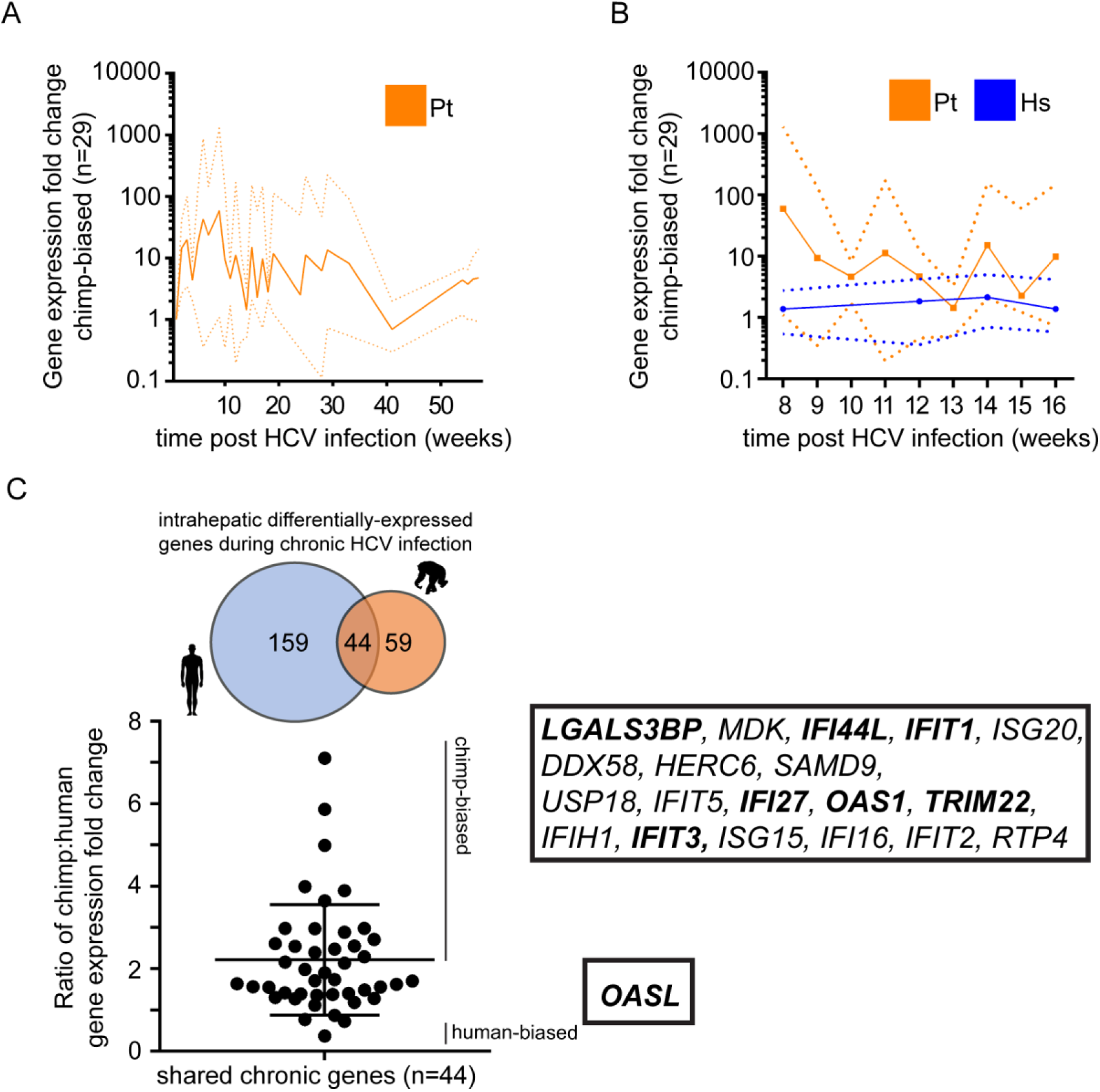
Comparative induction of hepatic transcripts during acute HCV infection in chimpanzees and humans. (A) Expression of ‘chimpanzee-biased’ differentially-expressed genes (n = 29) up to more than 50 weeks post infection. Chimpanzee-biased genes are shown as a combined mean (filled orange line) and range (dotted orange lines) of fold-change from all studies where any gene of the 29 genes was available over at most 1 year after initial infection. (B) Expression of differentially-expressed human and chimpanzee genes during 8 to 20 weeks post infection. Chimpanzee-biased genes are shown as a combined mean (filled orange line) and range (dotted orange lines) of fold-change from all 29 genes. The mean and range of gene expression for the human genes (blue filled and dotted lines respectively) equivalent to the chimpanzee-biased genes are shown. These data for chimpanzees are combined from different serial samples from different animals while human data is taken from a single biopsy from patients with an inferred time post infection. (C) Shared gene expression (overlapping 44 genes as shown in Venn diagram) during chronic infection in humans and chimpanzees as illustrated by a ratio of fold change in expression for humans and chimpanzees. Species biased genes (>2 fold enriched in either species) are listed to the side. All data are available in **Supplementary Data File 3.**

**S7 Fig.**
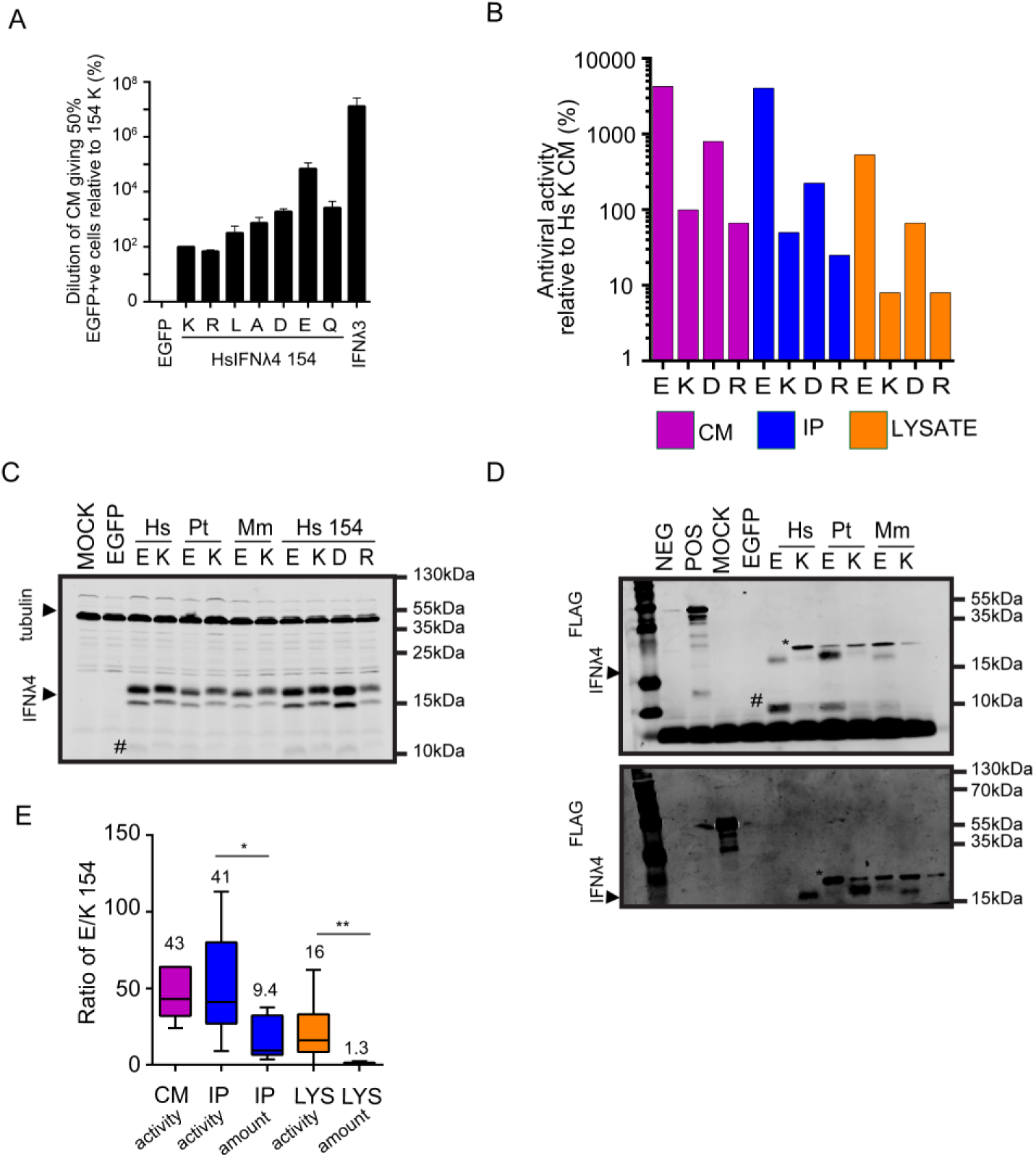
Mechanistic insight into variant E154. (A) Dilutions for inducing ∼50% EGFP positive cells for HsIFNλ4 IFNλ4 154 mutants (R, L, A, D, E and Q) using an EGFP-ISG15 reporter cell line assay relative to CM for HsIFNλ4 K154 (%). Data show mean +/- SEM from three experiments. (B) Antiviral activity of HsIFNλ4 isolated from CM (purple), intracellular lysate (blue) and immunoprecipitated CM (orange) for variants encoding E, K, R or D at position 154 in an anti-EMCV CPE assay relative to CM for wt HsIFNλ4 (K154 variant) in HepaRG cells. (C) Detection of intracellular IFNλ4 from different species as well as select mutants at position 154 (E, K, D and R) by Western blot analysis of lysates from plasmid-transfected producer HEK293T cells. The IFNλ4 variants were detected with an anti-FLAG antibody. Tubulin was used as a loading control. These samples were taken from the same experiment as in Fig 6E. A lower molecular weight product detected by the anti-FLAG antibody, which is potentially a degradation product, is highlighted with a ‘#’ (D) Alternative images from repeats of detection of extracellular IFNλ4 from different species by Western blot analysis of samples of FLAG-tag immunoprecipitated CM (1 ml) from plasmid-transfected producer HEK293T cells. A BAP-FLAG fusion protein was used an immunoprecipitation control (POS). IFNλ4 variants were detected with an anti-FLAG antibody. A lower molecular weight product detected by the anti-FLAG antibody, which is potentially a degradation product, is highlighted with a ‘#’. An upper band running near to the IFNλ4 species is shown (*) which is likely antibody fragments from the immunoprecipitation reaction. (E) Ratio of E154 to K154 for antiviral activity (all) or protein amounts (IP or LYS) of IFNλ4 found in CM, intracellular lysate and immunoprecipitated CM; where possible, data from the different species is combined. Data show median +/- minimum and maximum (n=6-8). Numerical data used for graph construction available in **Supplementary Data File 4 sheet 6.**

**S8 Fig.**
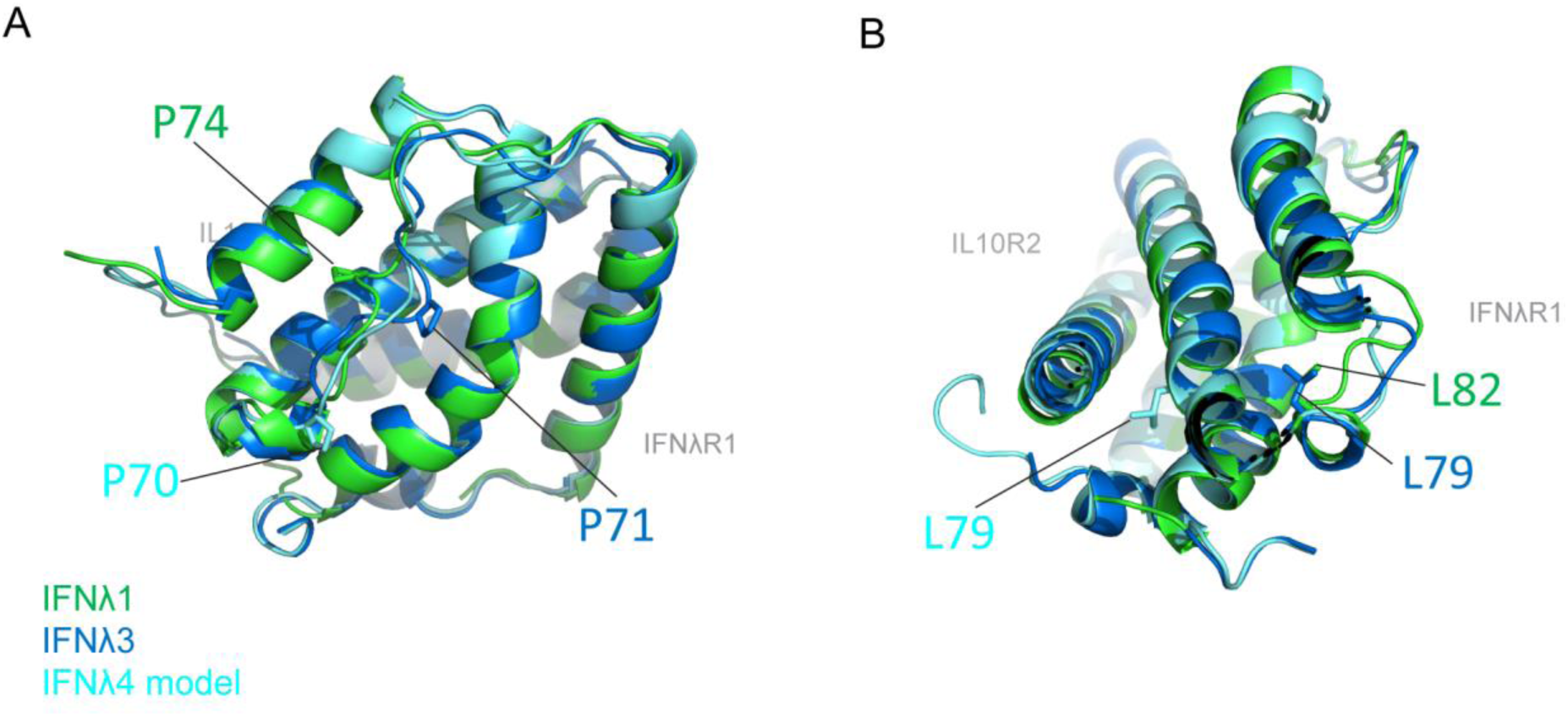
Structural modelling of P70 and L79 in HsIFNλ4. Modelled structures of HsIFNλ4 (light blue) are overlaid on the crystal structures for HsIFNλ1 (green) and HsIFNλ3 (dark blue). Panels A and B show respectively positions P70 and L79 in HsIFNλ4 and their homologous positions in HsIFNλ1 and HsIFNλ3 with reference to receptor subunit-binding interfaces (IFNλR1 and IL10R2 in grey). (A) P70 is found in a flexible proline-rich loop/helix facing the IFNλR1 receptor although no direct interactions between this region or receptor have been demonstrated. P70S might prevent required folding of the protein domain. (B) L79F is likely to disrupt packaging of the helices. Both P70 and L79 are highly conserved between IFNλ4 orthologues and paralogues (data not shown).

**S9 Fig.**
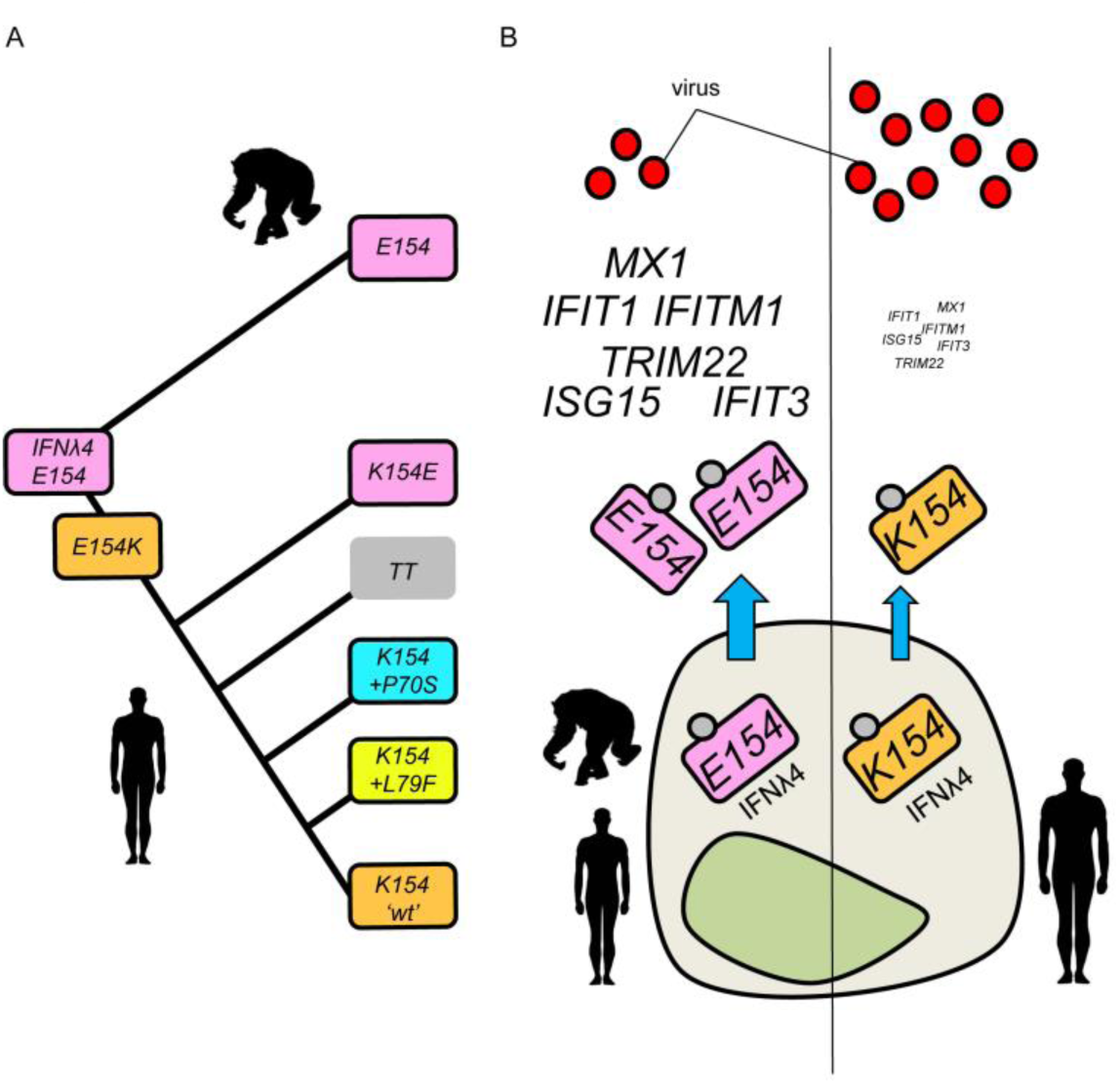
Evolution and functional impact of variation at position 154 of IFNλ4. (A) Inferred evolution of position 154 in humans and chimpanzees. The last common ancestor of humans and chimpanzees encoded the highly-conserved glutamic acid (E) at position 154 (purple). E154 was retained in chimpanzees but sequentially modified in the genus *Homo*, which includes humans. *Homo* IFNλ4 was first modified by substitution of E154 to lysine (K) (orange) and subsequent emergence of the frameshift TT allele (grey) or by the introduction of other substitutions (P70S [blue] or L79F [yellow]). IFNλ4 in humans with only the E154K change remains in the population and is considered wild-type (’wt’). (B) Impact of E and K encoded at position 154 in IFNλ4 on antiviral activity. Both IFNλ4 E154 (purple) and K154 (orange) are produced and glycosylated (grey circle) to similar levels inside the cell but IFNλ4 E154 is secreted more efficiently compared to IFNλ4 K154 (highlighted by blue arrows). Moreover, IFNλ4 E154 is also more potent than IFNλ4 K154. Subsequently, IFNλ4 E154 induces more robust interferon stimulated gene (ISG) expression (for example: *ISG15, IFIT1, MX1*) in target cells, leading to greater antiviral activity.

